# Reciprocal mechanochemical feedback couples neural crest migration and neurulation

**DOI:** 10.64898/2026.08.14.744800

**Authors:** Kai Weißenbruch, Lucas Alvizi, Jemima J. Burden, Roberto Mayor

## Abstract

Cephalic neurulation and neural crest migration are defining morphogenetic events of vertebrate head development. Although traditionally viewed as independent tissue-autonomous programs, their striking spatiotemporal overlap suggests functional coupling. Here, we show that these processes are linked by reciprocal mechanochemical feedback. We find that collective neural crest migration is essential for neural tube closure. As neural crest cells delaminate and invade the surrounding mesoderm, they remodel fibronectin at the neural crest-neural plate interface, creating a specialized extracellular matrix that both separates the two tissues and promotes neural tube morphogenesis by enabling radial intercalation and apical constriction of neural plate cells. This extracellular matrix remodeling requires neural crest-specific expression of the membrane-bound metalloproteinase MMP14. Conversely, neural tube morphogenesis drives neural crest migration. Mechanical compression generated during neural plate bending induces MMP14 expression in neural crest cells, triggering extracellular matrix remodeling that feeds back to facilitate neural tube closure. Together, our findings reveal that neurulation and neural crest migration are not independent morphogenetic programs but components of a self-reinforcing mechanochemical circuit that coordinates vertebrate head morphogenesis.

## Introduction

A central concept in developmental biology is that mechanical morphogenesis - the movement and shape changes of cells - follows chemical morphogenesis, which delivers the blueprint for tissue organization and enables cells to self-organize into higher-order three-dimensional structures^[1]^. However, growing evidence indicates that mechanical forces are not merely downstream consequences of biochemical patterning but also serve as instructive signals that coordinate tissue morphogenesis across scales through reciprocal feedback between mechanical forces and biochemical signalling^[2]^. While such reciprocal mechanochemical feedback networks have predominantly been studied in individual tissues or cell populations, developing organs comprise multiple neighboring tissues that undergo morphogenesis simultaneously. Whether these morphogenetic programs merely develop in parallel or instead exchange instructive mechanochemical cues to coordinate one another remains a fundamental question in biology.

Unlike chemical signals, which can arise and act in an autocrine manner within a homogeneous population of cells^[3]^, mechanical signaling requires force transmission between physically interacting structures. Consequently, instructive mechanical cues frequently emerge at interfaces between neighboring tissues and/or the extracellular matrix (ECM), given that such interfaces often possess significantly different physical characteristics^[4–6]^. Several studies suggest that mechanically coupled tissue interactions represent a widespread mechanism to coordinate morphogenesis in development and disease^[5, 7–18]^. However, the origin of these instructive mechanical cues and the mechanisms by which they are generated and exchanged between neighboring tissues in vivo remain poorly understood.

To explore how neighboring morphogenetic programs can mechanochemically coordinate one another, we use *Xenopus laevis* to investigate feedback during morphogenesis of the cephalic neuroectoderm. Cephalic neurulation, which drives neural tube (NT) formation, and collective migration of cephalic neural crest (NC) cells are two of the most prominent morphogenetic processes during early vertebrate head development, with devastating pathological consequences when disrupted^[19, 20]^. Before neurulation, the prospective NT and NC reside within the neuroepithelium as a genetically distinct but morphologically coherent epithelial tissue. With the onset of neurulation, however, both tissues rapidly diverge. The neural plate (NP) bends and converges towards the dorsal midline, whereas NC cells undergo epithelial-to-mesenchymal transition (EMT) and collectively migrate away from the NP^[21]^.

Extensive work has elucidated the molecular and mechanistic basis of both morphogenetic programs from a predominantly tissue-autonomous perspective. Cephalic NT morphogenesis is thought to be mainly driven by supracellular force patterns that induce coordinated apical constriction^[22, 23]^ to induce tissue bending, convergent extension along the midline^[24]^, as well as medial migration and cell intercalation^[25]^. Moreover, lateral geometric confinement has emerged as a key physical requirement for NT folding in minimal organoid systems^[26]^. Conversely, cephalic NC migration depends on collective chemotaxis and durotaxis together with synchronized supracellular contractions at the rear of the NC cluster, among several other intrinsic and extrinsic cues^[27]^.

Because these morphogenetic programs display striking tissue-autonomous characteristics and their movements proceed away from, rather than towards one another, the prevailing view is that NT morphogenesis and NC migration represent largely autonomous developmental programs. This view is further reinforced by trunk development, where NC delamination and migration begin only once NT closure has been completed^[21]^. In the prospective head, however, NT closure and NC migration occur simultaneously, with considerable spatiotemporal overlap across vertebrate species^[21]^. Whether this overlap merely reflects developmental coincidence or instead indicates functional and mechanical interdependence between the two tissues has remained unknown.

Here we show that collective cephalic NC migration is required for cephalic NT closure. As NC cells delaminate and invade the ECM secreted by the cephalic mesoderm, they remodel fibronectin (FN) along the NC-NP interface. This remodeled FN not only physically separates NC and NP but also enables NT morphogenesis by facilitating radial intercalation and apical constriction of NP cells during NT closure. Mechanistically, cephalic NC cells express the membrane-bound matrix metalloproteinase MMP14, which, together with contractility-driven retrograde FN treadmilling, mediates NC-dependent FN remodeling. Consequently, this remodeling relies on spatially localized MMP14 activity generated by collective migration rather than on diffusible protease activity. Strikingly, NC-specific MMP14 expression itself is mechanosensitive and is upregulated by compressive forces generated during neurulation, as neural plate bending progressively confines NC cells between the underlying mesoderm and the overlying ectoderm. Together, these findings reveal a reciprocal mechanochemical feedback loop in which NP bending mechanically activates NC invasion, whereas NC cells remodel the FN matrix to drive NT closure.

## Results

### FN is progressively deposited along the cephalic NC-NP interface and segregates both populations

To investigate the spatiotemporal development of cephalic NT closure and NC migration, we prepared serial transverse cryosections spanning the second through the third NC stream at stages before, during, and after NT closure (Supplement Figure 1a&b). Before neurulation, the prospective NT and NC reside within the neuroepithelium as a coherent epithelial tissue: a flat, multilayered NP (Sox2-positive), flanked by NC cells (Sox9-positive) on both sides (Figure 1a). With the onset of cephalic neurulation, NC and NP become progressively separated by a primary ECM, consisting of a thin FN layer that gradually wraps around the bending NP (Figure 1a-g and Supplement Figure 1a-f). We refer to this as FN at the NC-NP interface (Figure 1e&f).

**Figure 1:**
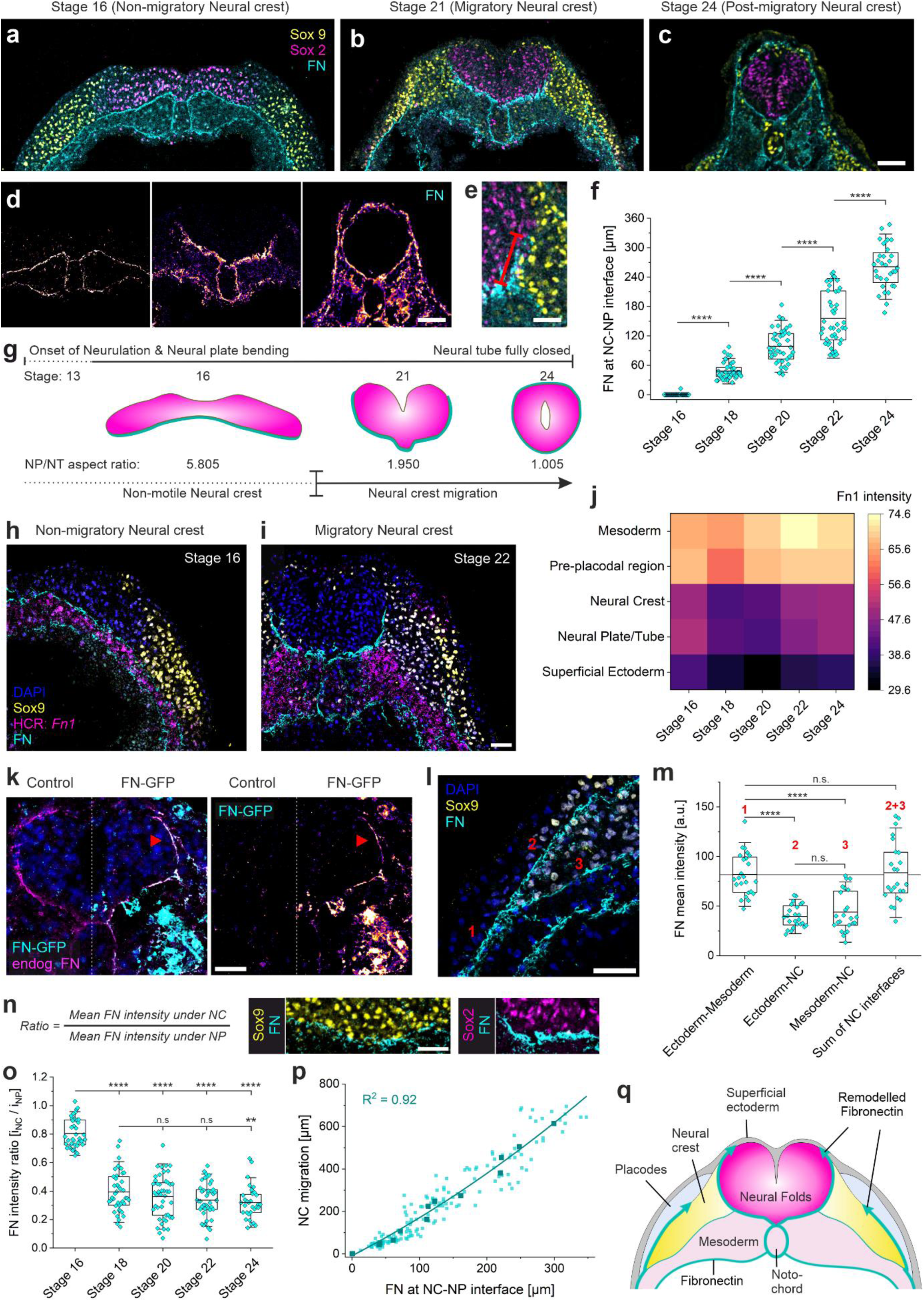
FN is progressively deposited along the cephalic NC-NP interface and segregates both populations. **a-c**, Transverse cryosections through the cephalic region of stage 16 (**a**), stage 21 (**b**), and stage 24 (**c**) embryos, stained for the NP marker Sox2 (magenta), the NC marker Sox9 (yellow), and FN (cyan). **d**, Insets showing FN from (**a**-**c**) as intensity LUT. **e**, Inset from **b**, highlighting where measurements were taken to quantify FN at NC-NP interface (red bar). **f**, FN length at NC-NP interface across stages. **g**, Illustration of NT morphogenesis (pink) with FN (cyan) progressively surrounding the forming NT. Timelines indicate NT morphogenesis (upper timeline) in relation to NC migration (lower timeline). **h&i**, Transverse cryosections of stage 16 (**h**) and stage 22 (**i**) embryos. HCR marking expression of *Fn1* (magenta), and immunostainings labeling NC cells (Sox9, yellow) and FN protein (cyan). Nuclei were labeled with DAPI. **j**, Heatmap of mean *Fn1* intensities across tissues and stages, measured by HCR signal in cryosections. **k**, Cryosection of the NT, showing co-localization of endogenous FN (magenta) and GFP-tagged FN (cyan and intensity LUT in right panel) along the NC-NP interface. FN-GFP was expressed from the mesoderm and GFP was additionally stained (anti-GFP Alexa Fluor 488) to enhance contrast. **l**, Inset showing leading edge of invading NC stream (Sox9, yellow), FN protein (cyan), and nuclei (DAPI). Numbers indicate different FN interfaces where measurements were taken. **m**, Mean FN protein intensity along indicated interfaces. **n**, Formula for ratio calculations displayed in (**o**). ROIs display FN interfaces underneath NC (Sox9, yellow) and NP (Sox2, magenta). **o**, Intensity ratio of FN protein underneath NC versus NP. **p**, Linear correlation of NC migration distance (quantified by the FN length at the NC-ectoderm interface) and length of FN at the NC-NP interface. Measurements were taken from cryosections of stage 16-24 embryos, as plotted in (**f**). **q**, Illustration depicting the emergence of FN along the NC-ectoderm and the NC-NP interface during cephalic neurulation and NC migration. Scale bars represent 100 µm in **a**-**d**, and 50 µm in **e**, **h**, **i**, **k**, **l**, **n**. For **f**, **m**, **o**, data are mean ± s.d. Statistical analysis was performed using Dunnett’s test (**f** and **o**) and Tukey’s test (**m**); n.s. *P* > 0.05; \*\**P* ≤ 0.01; \*\*\*\**P* ≤ 0.0001. *n* = 32-45 measurements from 10 different embryos (**f**, **o**, **p**), *n* = up to 45 measurements from 10 different embryos (**j**), and *n* = 26 measurements from 12 different embryos (**m**).

Transcription of FN was analyzed by HCR, showing that it was expressed neither by NP nor NC cells, but instead by the underlying mesoderm and the pre-placodal region anterior to the eye vesicles (Figure 1h-j and Supplement Figure 2a-g). FN expression was detected throughout the mesoderm between stage 16 and 24 (Figure 1j), before becoming progressively restricted to placodal regions from stage 25 onwards (Supplement Figure 2d). As FN is not expressed by the NP or NC, we asked where the FN deposited at the NC–NP interface originates. To identify the origin of this FN, we blocked its expression specifically in the mesoderm or in the pre-placodal region in one half of the embryo. Depleting FN from the pre-placodal source had no significant effects on FN distribution around the cephalic NC-NP interface (Supplement Figure 3a&c) and little effect on NC migration (Supplement Figure 3e&h). By contrast, depletion of mesoderm-derived FN resulted in an almost complete loss of FN throughout the targeted side of the embryo, including the regions surrounding the cephalic NC and NP (Supplement Figure 3b&d). Under such conditions, NC migration was severely disrupted (Supplement Figure 3f&h).

Together, these findings indicate that the FN matrix separating NC and NP during cephalic neurulation is generated by remodeling pre-existing mesoderm-derived FN rather than by *de novo* synthesis. Consistent with this interpretation, expression of GFP-tagged FN in the mesoderm reconstituted the endogenous localization of FN (Supplement Figure 2h&i) and resulted in the incorporation of GFP-positive fibrils into the endogenous FN at both the NC-NP interface (Figure 1k) and the NC-ectoderm interface (Supplement Figure 2j&k).

### NC migration spatiotemporally correlates with FN deposition at the NC-NP interface

To understand how FN is relocalized from the mesoderm to the NC-NP interface, we next investigated which cell population drives this process. Because GFP-positive FN fibrils were incorporated not only into the NC-NP interface but also into the NC-ectoderm interface (Figure 1l&m and Supplement Figure 2j&k), we hypothesized that cephalic NC cells actively remodel FN at both sides. Consistent with this hypothesis, cephalic NC cells invade the FN meshwork at the onset of migration, splitting the continuous ECM into two layers: one remaining at the NC-mesoderm interface and the other covering the NC-ectoderm interface (Figure 1l and Supplement Figure 1g&h). The combined FN intensity of these two newly formed interfaces equaled the intensity of the FN layer ahead of the migrating NC cluster, suggesting that the existing ECM is split rather than newly synthesized (Figure 1m). Concurrently, a second FN cleft formed along the NC-NP interface at the rear of the NC cluster, which subsequently developed into the FN at the NC-NP interface (Supplement Figure 1h). We quantified the FN distribution along the NC-NP interface throughout cephalic neurulation to determine whether this FN remodeling is also coordinated with NC migration, and found both events were spatiotemporally correlated. The first FN along the NC-NP interface appeared at the onset of NC delamination and expanded linearly from stage 18 to stage 24, matching the timing of NC migration (Figure 1f&g). As the FN expanded along the NC-NP interface, FN intensity progressively decreased at the NC-mesoderm interface, while FN levels at the NP-mesoderm interface remained unchanged (Figure 1n&o and Supplement Figure 1h&i). Consequently, the length of the FN along the NC-NP interface correlated linearly with the length of the NC streams, indicating that FN redistribution is temporally coupled to NC migration (Figure 1p).

Together, these data suggest that cephalic NC cells remodel FN to localize it at the NC-NP interface, driving tissue segregation during cephalic neurulation (Figure 1q and Supplement Figure 1b).

### Cephalic NC cells express MMP14 to remodel FN

To investigate the mechanism underlying NC-dependent FN remodeling, we focused on matrix metalloproteinases (ADAMs and MMPs) as likely candidates. Cephalic NC cells in *Xenopus laevis* express at least three MMPs/ADAMs (Supplement Figure 4a). ADAM13 is specifically expressed in the cephalic NC^[28]^ but has been implicated in NC induction^[29]^, transcriptional regulation^[30]^ and Cadherin-11 shedding^[31, 32]^, rather than directly promoting NC migration^[30, 33]^. MMP2 is broadly expressed in the cephalic mesenchyme^[34]^ and is detected in NC cells only from stage 24 onwards, when cephalic NC migration is almost finished^[35]^. Consistent with this, we detected no NC-specific MMP2 expression at earlier stages (Supplement Figure 4a), and previous studies showed that MMP2 inhibition had no cell autonomous effect on NC migration in *Xenopus laevis^[36]^*. By contrast, MMP14 is a membrane-anchored MMP expressed on cephalic NC cells but not the NP (Figure 2a&b and Supplement Figure 4a). Moreover, depletion of MMP14 blocked cephalic NC migration^[36]^ (Supplement Figure 4b&c), making it a likely candidate.

**Figure 2:**
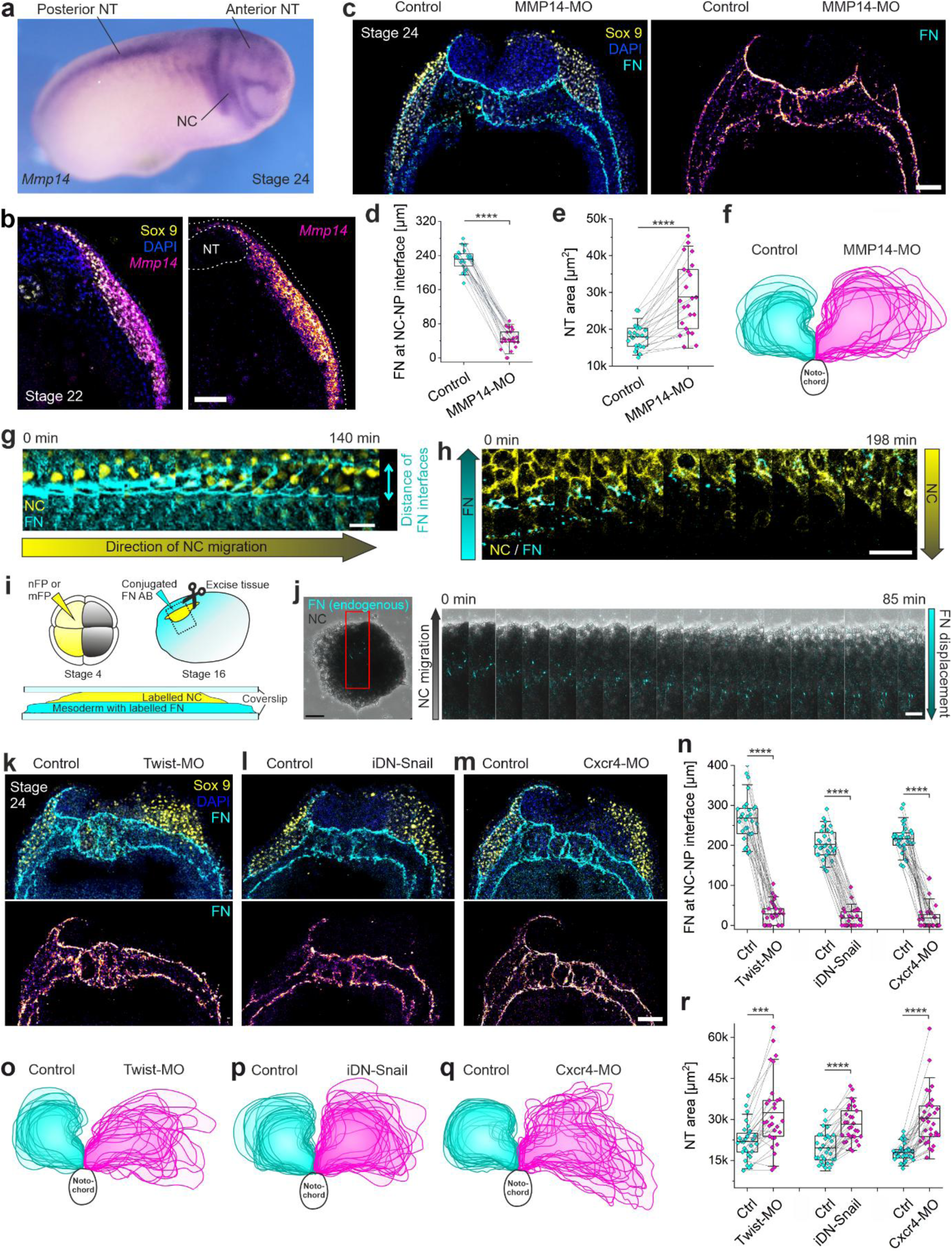
NC migration-dependent FN remodeling through MMP14 is a prerequisite for NT closure. **a**, Stage 24 embryo stained by colorimetric ISH for *Mmp14*. **b**, Transverse section of stage 22 embryo. Fluorescent ISH marking expression of *Mmp14* (magenta and LUT in right panel), and immunostaining labeling NC cells (Sox9, yellow) and nuclei (DAPI). **c**, Transverse section of stage 24 embryo, in which MMP14 expression was blocked in the right side of the embryo. Staining against the NC marker Sox9 (yellow), FN (cyan and LUT in right panel) and nuclei (DAPI). **d**, FN length at NC-NP interface. **e**, Cross-sectional areas of NT hemispheres. **f**, Overlayed areal projections of NT hemispheres in control (cyan) and MMP14-depleted (magenta) embryo halves from 18 sections and 12 embryos. **g**, Time-lapse montage of nGFP expressing NC cells invading mesoderm-derived FN (labeled with Alexa Fluor 555-conjugated antibody), corresponding to Supplement Video 2. Yellow arrows indicates direction of NC migration. **h**, Time-lapse montage of mGFP expressing NC cells retrogradely displacing mesoderm-derived FN (labeled with Alexa Fluor 555-conjugated antibody), corresponding to Supplement Video 3. Cyan and yellow arrows indicate FN displacement and direction of NC migration, respectively. **i**, Illustration of the assay shown in (**g**, **h**). **j**, Time-lapse montage of NC explant from embryo in which mesoderm-derived FN was labelled with Alexa Fluor 555-conjugated antibody. Explant was seeded on FN-coated dish and FN displacement was monitored in relation to the registered leading edge of the NC cluster. **k-m**, Transverse sections of stage 24 embryos, in which the right side of the embryo was injected with a *Twist1* morpholino (**k**), inducible dominant negative Snail (**l**), or *Cxcr4* morpholino (**m**). Sections were stained for the NC marker Sox9 (yellow), FN (cyan and LUT in lower panels), and nuclei (DAPI). **n**, FN length at NC-NP interface, quantified in control and injected embryo halves of respective embryos. **o**-**q**, Overlayed areal projections of control (cyan) and injected (magenta) NT hemispheres from 18 Twist-MO, 19 iDN-Snail, and 19 Cxcr4-MO sections and 10 embryos each. **r**, Cross-sectional areas of NT hemispheres. Scale bars represent 100 µm in **b**, **c**, **k**, **l**, **m**, 50 µm in **h**, **j** (overview), and 20 µm in **g**, **j** (montage). For **d**, **e**, **n**, **r**, data represent mean ± s.d. Statistical analysis was performed using two-tailed Mann-Whitney test (**n**) and two-tailed *t*-test (**d**, **e**, **r**). \*\*\**P* ≤ 0.001; \*\*\*\**P* ≤ 0.0001. *n* = 23 measurements from 12 different embryos (**d** and **e**); *n* = 29, 30, and 32 measurements from 10 different embryos, each (**n** and **r**).

Analysis of transverse sections from MMP14 morphants revealed a marked reduction in FN at the NC-ectoderm and the NC-NP interface (Figure 2c&d). Although small FN clefts still formed at both interfaces, suggesting that the onset of NC migration is not affected, further progression of migration was impaired (Figure 2c and Supplement Figure 4f). Consistent with NC cells redistributing mesoderm-derived FN, FN intensity remained significantly higher beneath the NC following MMP14 depletion, whereas no difference was observed beneath the NP (Supplement Figure 4f-h). To determine whether MMP14 directly regulates NC cell motility, we examined the behavior of MMP14-deficient NC cells ex vivo. When cephalic NC explants were cultured on FN-coated coverslips, both control and MMP14-depleted cells were motile and dispersed randomly from the cluster (Supplement Figure 4i&j and Supplement Video 1), with individual MMP14-depleted NC cells migrating with comparable directionality and even higher speed than control cells (Supplement Figure 4k&l).

These findings indicate that MMP14 is dispensable for intrinsic NC motility but required for FN cleavage during migration within the confined three-dimensional environment in vivo. While NC cells migrate dorsolaterally, they split the pre-existing FN matrix at the leading edge of the NC cluster into two. A similar splitting event at the rear of the NC cluster can initiate FN remodeling at the NC-NP interface. However, this requires that NC cells not only cleave but in addition also retrogradely displace FN opposite to their direction of migration. To test this idea, we fluorescently labelled cephalic NC cells and explanted them at the premigratory stage together with the underlying mesoderm, in which FN fibrils had been labelled by subcutaneous injection of an Alexa Fluor 555-conjugated FN antibody. This explant was then sandwiched between two FN-coated coverslips and imaged by time-lapse microscopy (Figure 2i). Upon NC migration, we observed FN splitting events, where large volumes of NC cells invaded into FN-rich regions (Figure 2g and Supplement Video 2). In addition, we observed retrograde displacement of labeled FN fragments against the direction of leader cells (Figure 2h and Supplement Video 3). To determine whether NC cells can mediate retrograde FN transport independently of mesoderm, we again labelled endogenous FN and explanted NC clusters at stage 20, after NC cells had already begun to migrate into the endogenous FN. As small fragments of labelled endogenous FN remained associated with the NC, we were able to monitor their displacement in time-lapse series ex vivo. Consistent with our hypothesis, NC cells transported FN fragments towards the center and rear of the cluster during migration (Figure 2j).

In sum, our data indicate that migrating NC cells use MMP14 to cleave FN and retrogradely translocate it towards the NC-NP interface.

### NC-dependent FN remodeling couples NC migration and NT morphogenesis

Strikingly, depletion of MMP14 not only inhibited NC migration and FN remodeling but also led to significant neurulation defects in the cephalic area. Upon MMP14 depletion, NT cross-sectional areas were increased compared to the contralateral control side (Figure 2e) and NTs failed to close (Figure 2f and Supplement Figure 4d&e). As MMP14 is a membrane-anchored MMP, its FN remodeling capacity is intrinsically linked to NC movement. We therefore hypothesized that NC migration and NT morphogenesis are functionally coupled through NC migration-dependent FN remodeling.

To explore this hypothesis, we first checked if NC migration by itself is necessary for NT closure. We specifically blocked cephalic NC migration using multiple independent approaches. An antisense morpholino against *Cxcr4* inhibits collective NC chemotaxis^[37]^, whereas a *Twist1* morpholino or an inducible dominant-negative Snail construct (iDN Snail) blocks NC EMT and delamination^[38, 39]^ (Supplement Figure 5a-e). Suppressing NC migration abolished FN redistribution along the NC-NP interface in all cases (Figure 2k-n). Moreover, the NT consistently failed to close at the midline (Figure 2k-q and Supplement Figure 5f-j) and exhibited significantly larger cross-sectional areas than the contralateral control side (Figure 2r), showing that NC migration is required for cephalic NT closure. This functional coupling was restricted to the cephalic region, as NT morphogenesis in the trunk remained unaffected (Supplement Figure 5k), consistent with reports showing that trunk NT closure is completed before NC cells breach the surrounding ECM and migrate away from the NT^[21]^.

As inhibition of NC migration impaired both FN redistribution at the NC-NP interface and NT closure, we next asked if NC migration promotes NT closure specifically through FN remodeling. Therefore, we ectopically expressed FN as mosaic in the superficial ectoderm of control embryos and FN remodeling-deficient MMP14 morphants to test if restoring FN at the NP-NC interface could rescue NT closure in the absence of NC-mediated FN remodeling (Supplement Figure 6). Ectopic expression of FN from the superficial ectoderm in control embryos had mixed effects on NC migration, with most embryos showing reduced NC stream length (Supplement Figure 6a&d), while NT closure was only mildly affected (Supplement Figure 6e,h,k). Strikingly, ectopic expression of FN from the superficial ectoderm in FN-remodeling deficient MMP14 morphants did not rescue NC migration (Supplement Figure 6b-d) but significantly improved NT closure (Supplement Figure 6f-k).

Together, these data suggest that NC-mediated FN remodeling functionally couples collective NC migration and NT morphogenesis.

### FN shapes NT morphogenesis

Following these results, we explored the role of FN during NT morphogenesis in more detail. One of the major mechanisms driving NT morphogenesis in the cephalic region is apical constriction^[22]^. We therefore asked whether apical constriction was affected when MMP14 activity and thus FN remodeling by the cephalic NC was blocked in one half of the embryo. Live cell imaging of whole mount labelled embryos showed that apical constriction was significantly slowed down and eventually halted in the targeted half compared with the control side (Figure 3a-c and Supplement Video 4).

**Figure 3:**
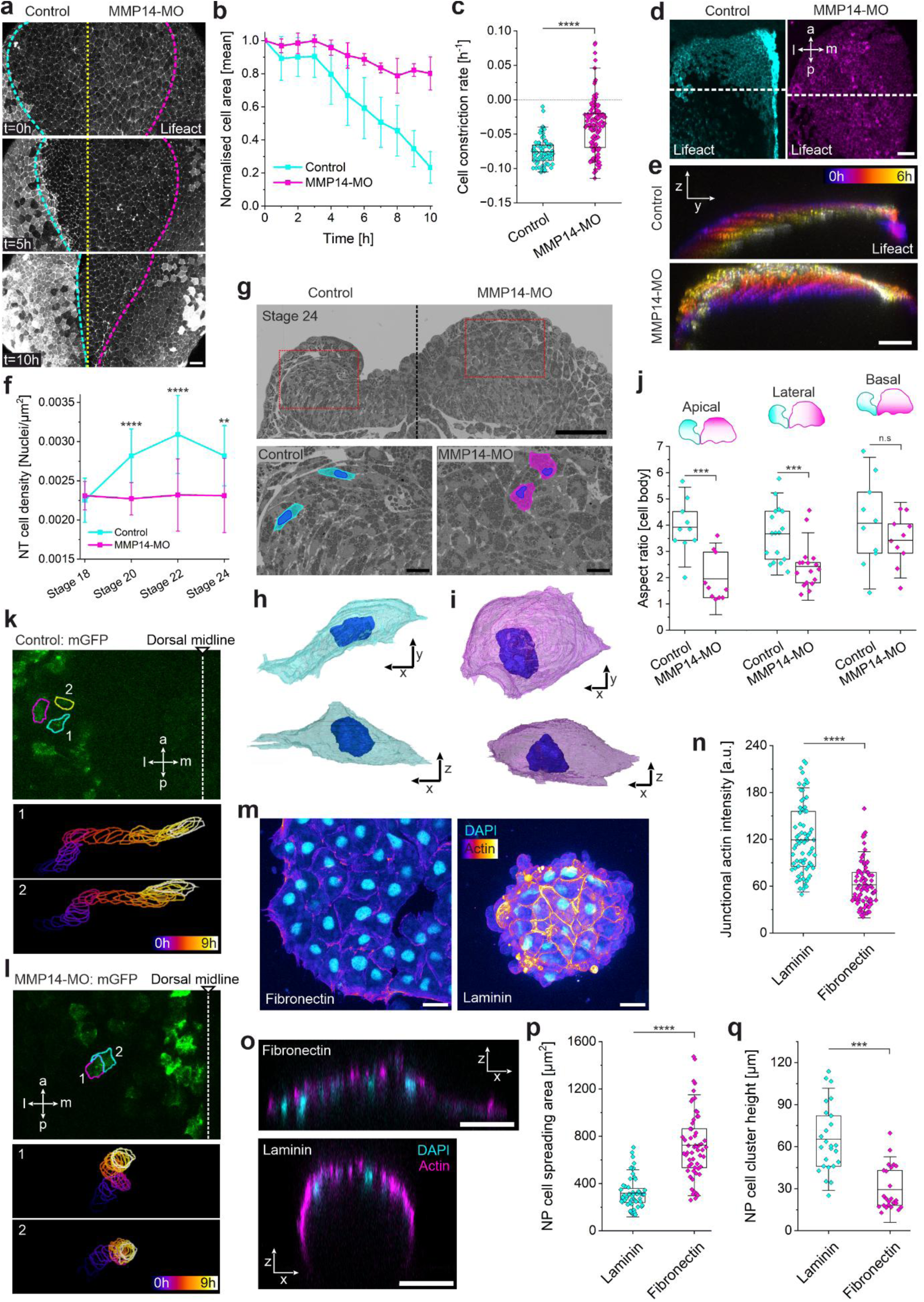
FN facilitates cell polarization for radial intercalation of basolateral NP cells. **a**, Time-lapse montage (dorsal view) of embryo expressing Lifeact-RFP and MMP14 depletion in right side of the embryo (Supplement Video 4). Dashed line marks midline (yellow), lateral NT edge in control (cyan), and lateral NT edge in MMP14-depleted side (magenta). **b**, Mean change of NT cell apical area over 10 h in control and MMP14-depleted embryo half. **c**, Mean constriction rate of individual NT cells in control or MMP14-depleted embryo halves. **d**, Cephalic region of left embryo half from control or MMP14-depleted embryos expressing Lifeact-RFP. Dorsal midline is to the right; white dashed lines indicate resliced ROIs. **e**, Temporally color-coded orthogonal projection of control or MMP14-depleted cephalic NT area (Dorsal midline to the right), corresponding to Supplement Video 5. **f**, Cell density in NT hemispheres of control and MMP14-depleted embryos during stages 18-24. **g**, Single-plane electron micrograph from a transverse section of stage 24 embryo with MMP14 depletion in right embryo half. Lower panels show ROIs marked by red boxes. Individual NT cells and nuclei were pseudocolored to highlight shapes. **h**-**i**, Reconstructed 3D cell and nucleus shape of a NT cell from control (**h**) and MMP14-depleted embryo half, corresponding to Supplement Video 6 and 7. **j**, Aspect ratios of NT cell cross-sections in ROIs depicted in legend. **k**-**l**, Still image (t = 0 h) and color-coded temporal projection of lateral NT cell shapes in control (**k**) and MMP14-depleted (**l**) embryo half during NT closure (Supplement Video 8). Numbers indicate which cells were followed. **m**, Maximum projection of NP explant on FN (left panel) or LN coating (right panel). Nuclei (DAPI) and actin cytoskeleton (phalloidin, LUT) were labeled. **n**, Junctional actin intensity in NP explants on FN or LN coating. **o**, Orthogonal projections of NP explants on FN (top) or LN (bottom). Nuclei (DAPI) and actin cytoskeleton (phalloidin, magenta) were labeled. **p**-**q**, Mean NP cell spreading area (**p**) and cluster height (**q**) on FN or LN. Scale bars are 100 µm in **d**, **e**, **g**; 50 µm in **a**, **o**; 20 µm in **m**, **g** (insets); and 5 µm each in **h**, **i**. For **b**, **c**, **f**, **j**, **n**, **p**, **q**, data represent mean ± s.d. Statistical analysis was performed using Tukey’s test (**f**), two-tailed Mann-Whitney test (**c**, **n**, **p**, **q**), and two-tailed *t*-test (**j**). n.s. *P* > 0.05; \*\**P* ≤ 0.01; \*\*\**P* ≤ 0.001; \*\*\*\**P* ≤ 0.0001. *n* = 4 control and 5 MMP14-MO embryos with 300-400 individual cells (**b**); *n* = 77 control and 134 MMP14-MO cells from 3 embryos each (**c**); *n* = 10, 14, 19, and 16 sections from 10-12 embryos each (**f**); *n* = 10, 17, and 10 cells in single SEM slice (**j**); *n* = 74 (LN) and 86 (FN) junctions (**n**), and *n* = 60 (LN) and 58 (FN) cells (**p**) from 20 explants each; *n* = 25 explants for LN and FN each (**q**).

Although apical constriction was impaired, several observations argue against FN directly regulating this process. Apical constriction begins before the onset of MMP14 expression and migration of the NC^[21]^. Moreover, the machinery driving apical constriction in the NP remained in place and active, even when MMP14 activity was blocked (Supplement Figure 7a-c). Finally, apical constriction occurs primarily in the most superficial neuroepithelial cell layer^[40]^, whereas only basolateral cells of the multilayered *Xenopus* neuroepithelium are in direct contact with FN. Together, these observations suggest that the effect of impaired FN remodeling on apical constriction is indirect. We therefore asked whether FN instead regulates behavior of basolateral cells during NT morphogenesis.

Because of the initially multilayered architecture of the NP in *Xenopus*, NT cells in the cephalic region were shown to undergo radial intercalation during NT closure^[25, 41]^. These intercalation events correlate with the timing of NC delamination and migration, and therefore with the emergence of FN at the basolateral side of the NT. Thus, we reasoned that FN could promote basolateral cell shape changes and radial intercalation within the NT. Such cell shape changes could facilitate basal relaxation and thereby act in synergy with apical constriction of superficial cells^[42–44]^.

To test this hypothesis, we first monitored the morphogenesis of the NT over time. Orthogonal projections revealed that the NT in control embryos shitfs towards the midline, while it bulges significantly out of plane and extends dorsolateral in MMP14 morphants (Figure 3d&e and Supplement Video 5). When we calculated the cell density of NT halves in control embryos and MMP14 morphants, we found a significant increase in density until stage 22 in the control, whereas the density remained constantly low in MMP14-depleted embryos (Figure 3f). This is consistent with the increased NT hemisphere area observed when NC migration was blocked (Figure 2e&r) and suggests an intercalation defect in the NT in the absence of FN.

To get a more direct analysis of cell intercalation, we next used volumetric scanning electron microscopy (array tomography^[45]^) to analyze the cellular morphology of individual basolateral NT cells in control and MMP14 morphant embryos. Intercalating cells were shown to possess a spindle-like shape with elongated nuclei, a characteristic of motile cells that actively reposition themselves in relation to their neighbors^[46, 47]^. In line with this, basolateral NT cells in control embryos possess a spindle-like morphology with elongated nuclei (Figure 3g&h and Supplement Video 6), marked by high aspect ratios of the cell body and nucleus (Figure 3j and Supplement Figure 7d). In MMP14-depleted embryos, basolateral NT cells showed a cuboid, unpolarized morphology with rounder nuclei (Figure 3g&i and Supplement Video 7), while only very basal cells that were in contact with the mesoderm-derived FN showed an elongated morphology (Figure 3j and Supplement Figure 7d). This observation suggests that FN is required to polarize NP cells, as apical and lateral cells lack contact with FN in MMP-14-depleted embryos due to the absence of FN at the NC-NP interface. Consequently, only basal cells remain in contact with FN, and these are the only cells that exhibit a polarized phenotype similar to control cells.

Time-lapse recordings of individual NT cells in control and MMP14-depleted embryos during neurulation suggest that these different cell phenotypes correlated with their ability to intercalate (Figure 3k&l). While elongated control cells in WT embryos translocated mediolaterally towards the midline (Figure 3k and Supplement Video 8), cuboid NT cells in MMP14 morphants failed to undergo these mediolateral translocations and instead remained at their initial position or translocated along the anterior-posterior axis (Figure 3l and Supplement Video 8).

Finally, we tested if FN alone is sufficient to induce cell elongation and intercalation. In addition to FN, Laminin (LN) is expressed around the Notochord and the adjacent mesoderm from stage 15 in the trunk and from stage 19 in the cephalic region (Supplement Figure 8a-j). However, LN was absent above the hinge region, while FN surrounded the whole NT (Supplement Figure 8k-n), making it the sole ECM component facing apical and lateral NT cells. We therefore cultivated NT explants ex vivo on FN or LN, respectively. On LN, NT cells remained in a compact cluster with cuboid shapes (Figure 3m), dense cell-cell contacts (Figure 3n), and low de-wetting^[48]^ (Figure 3o-q), in line with findings in other cell culture systems^[49]^. On FN, NT cells loosened individual junctions (Figure 3m&n) and spread towards a flatter and more elongated shape (Figure 3o-q). Moreover, when we placed two explants in proximity, we observed intercalation events on FN but not on LN (Supplement Figure 7e-g and Supplement Video 9). Together, these findings show that FN is sufficient to convert NT cells from a cuboid, tightly interconnected state into a polarized spindle-shaped phenotype that enables mediolateral intercalation into neighboring rows, similar to observations reported in other tissues and model organisms^[7, 50–54]^.

### NT closure confines mechanosensitive NC cells

Having established that migrating NC cells shape NT closure through FN remodeling, we next asked whether NT closure reciprocally regulates NC migration. To test this, we inhibited NT closure by using a morpholino against Shroom3, which is known to inhibit apical constriction in the cephalic NP^[22, 23]^ (Figure 4a). At the same time, embryos with open NTs also showed impaired NC migration, with very short NC streams (Figure 4b&c) and consequently missing FN deposition at both the NP-NC interface and the NC-ectoderm interface (Figure 4d&e).

**Figure 4:**
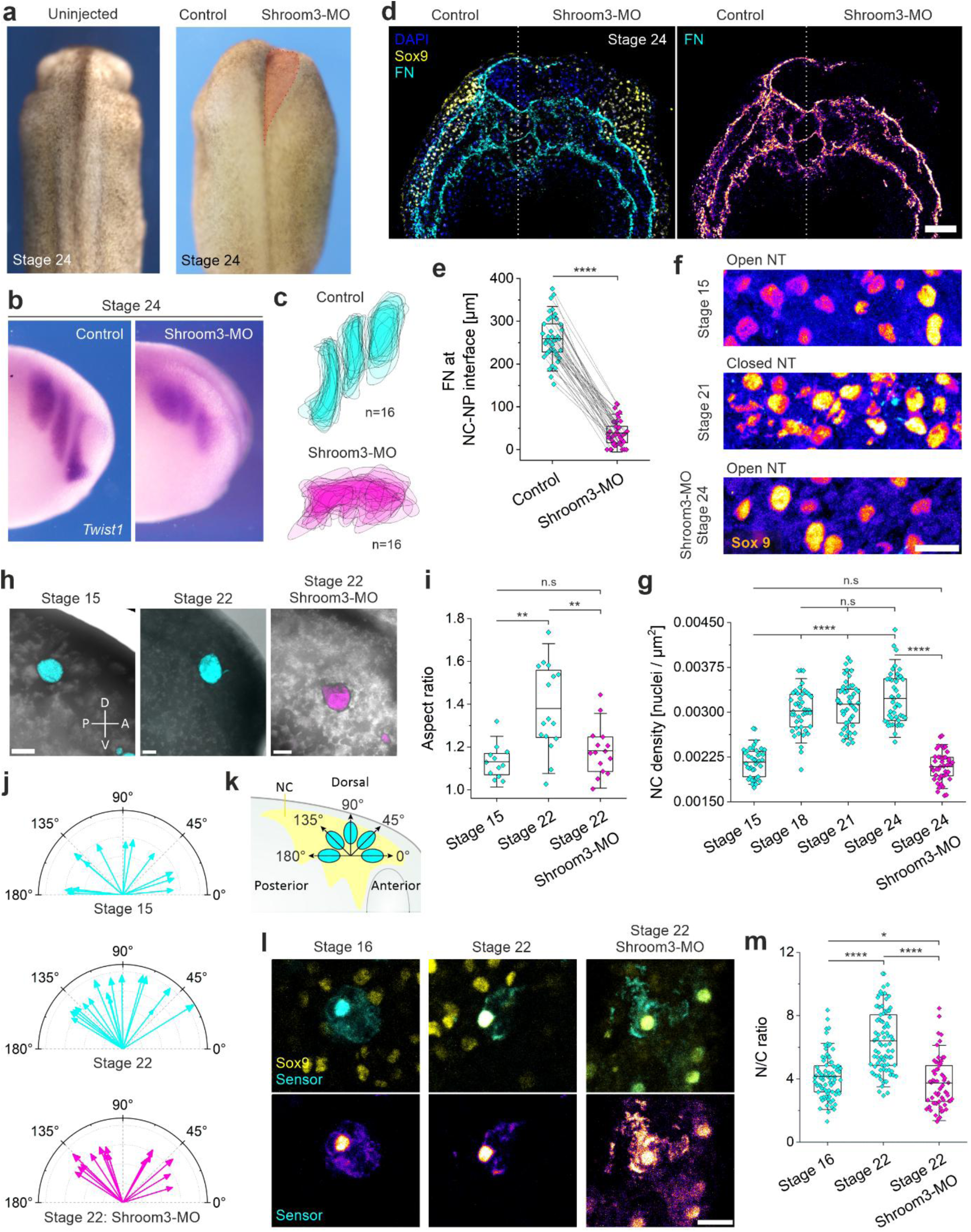
NP bending compresses mechanosensitive cephalic NC cells. **a**, Dorsal view of uninjected (left panel) and Shroom3-depleted (right side) embryo at stage 24. Highlighted area in red marks where NT failed to close. **b**, Embryos were injected with *Shroom3* morpholino in one side and stained for the NC marker *Twist1* by ISH. **c**, Overlayed areal projections of NC streams in control (cyan) and Shroom3-depleted (magenta) embryo halves. **d**, Transverse section of stage 24 embryo, in which Shroom3 expression was blocked in right side of the embryo. Staining against the NC marker Sox9 (yellow), FN (cyan and LUT in right panel) and nuclei (DAPI). **e**, FN length at NC-NP interface. **f**, Close-up of NC streams before NT closure (stage 15, top panel), after NT closure (stage 21, middle panel), and when NT closure was blocked by *Shroom3* morpholino injection (stage 24, lower panel). NC nuclei were marked by Sox9 staining (intensity LUT). **g**, NC density, plotted as number of nuclei per µm^2^, across different developmental stages (before, during, after NT closure and when NT closure was blocked). **h**, Fluorescent oil droplets were injected into the cephalic NC area before NT closure (stage 15, left panel), after NT closure (stage 22, middle panel), and when NT closure was blocked by *Shroom3* morpholino injection (right panel). **i**, Oil droplet elongation measured as aspect ratios). **j**, Compass plots of oil droplet elongation angles before NT closure (stage 15, top plot), after NT closure (stage 22, middle plot), and when NT closure was blocked (stage 22: Shroom3-MO, lower plot). Angles indicate droplet elongation direction in respect to embryo axis, and arrow length indicates elongation ratio itself. **k**, Schematic of droplet elongation angles. **l**, Embryos at depicted stages were injected with nuclear mechanosensor construct (L_NLS 41 kDa) alone or together with *Shroom3* morpholino to block NT closure. Sections were stained for Sox9 (yellow) to distinguish NC cells. **m**, Quantification of nuclear-to-cytoplasmic (N/C) ratio of the sensor. Scale bars represent 100 µm (**d**), 50 µm (**h**), 20 µm (**f**), 10 µm (**l**). For **e**, **g**, **i**, **m**, data represent mean ± s.d. Statistical analysis was performed using two-tailed *t*-test (**e**), Tukey’s test (**g**), and Dunn’s test (**I** and **m**). n.s. *P* > 0.05; \**P* ≤ 0.05; \*\**P* ≤ 0.01; \*\*\*\**P* ≤ 0.0001. *n* = 16 embryos (**c**); *n* = 43 sections from 12 different embryos (**e**); *n* = 12, 16, and 15 embryos, respectively (**i**); *n* = 41, 44, 49, 43, 44 sections from 12 different embryos (**g**); *n* = 81, 87, 62 sections from 12 different embryos (**m**).

Given that FN remodeling is functionally initiated by the NC rather than by the NP, we reasoned that NT closure impairs NC migration and FN remodeling in a non-cell autonomous manner. Transplantation experiments confirmed this, showing that NC cells cannot migrate when grafted into a Shroom3-depleted host, while Shroom3-depleted NC cells migrated when grafted into a control host (Supplement Figure 9a&b). Furthermore, ex vivo, both control and Shroom3 morphant NC cells were motile (Supplement Video 10). Single cells migrated with comparable directionality, while migration speed of single cells even increased in Shroom3-depleted explants (Supplement Figure 9c-f).

A potential explanation comes from the emerging link between cell compression and invasive properties. For example, melanoma cells become more invasive upon compression^[55]^. Due to the similarities between cancer cell invasion and NC migration^[56]^, we speculate that NC cells might be compressed between the mesoderm and ectoderm upon bending of the NP and subsequent closure of the NT. As the cephalic NC has been shown to respond very sensitively to mechanical stimuli during their induction^[57, 58]^, EMT^[14, 15]^, and migration^[16]^, we reasoned that such compression might trigger invasive characteristics in the cephalic NC by acting as a stimulus for NC EMT and FN remodeling.

To test this hypothesis, we first confirmed an increase in NC cell density with progressing NT closure in vivo. NC density increased from non-migratory to pre-migratory and migratory stage but remained low, when we blocked Shroom3 to inhibit NT closure (Figure 4f&g and Supplement Figure 10a&b). To further test the idea that cells within the NC tissue were under compression, we analyzed the deformability of oil droplets^[59, 60]^ injected into the extracellular space of the NC tissue and compared their deformability before and after NT closure. Droplets remained more round before NT closure and were not preferentially elongated along any specific embryonic axis (Figure 4h-k). After NT closure, however, droplets were more elongated and oriented with their longest axis mainly along the dorsoventral embryo axis (Figure 4h-k). Again, these effects were not pronounced when NT closure was blocked, as droplets in Shroom3-depleted embryos had lower aspect ratios and the elongation angle was oriented more along the anterior-posterior axis (Figure 4h-k).

To further test if NC cells are responsive to this compression, we made use of a previously published nuclear mechanosensor (L_NLS 41 kDa)^[61]^, whose influx into the nucleus from the cytosol increases in response to mechanical compression of the nucleus and the dilation of its nuclear pore complexes^[62]^. To monitor individual NC cells, we expressed the sensor as a mosaic in the neuroectoderm and co-stained Sox9 in cryosections to distinguish NC from other cell types (Figure 4l). Quantifying the sensors’ nuclear-to-cytoplasmic ratio (N/C ratio) in stages before and after NT closure revealed a significant increase in N/C ratio after closure of the NT (Figure 4l&m). Once again, blocking NT closure by morpholino injection against Shroom3 decreased the nuclear fraction of the sensor, showing a comparable mean N/C ratio to control embryos before NT closure (Figure 4l&m).

Together, these findings indicate that bending of the NP and subsequent NT closure mechanically compresses cephalic NC cells and that this compression is sensed by the NC.

### NC compression arises from a combination of mesodermal stiffening and tension increase in the superficial ectoderm

We next asked how mechanical compression of the NC arises from the surrounding tissues. It has been previously shown that the head mesoderm underneath the NC stiffens due to convergent extension and that this triggers the onset of NC EMT and delamination^[15]^. This increase in stiffness occurs during the same developmental stages as NP bending. In addition, we noticed that the superficial ectoderm was bulged outward above the NC in Shroom3 morphants (Figure 4d), suggesting that tension in the superficial ectoderm might also be affected. To confirm this, we quantified ectodermal tension above the NC by measuring its recoil velocity via laser ablation (Figure 5a&b, Supplement Figure 10c-e, and Supplement Video 11). In control embryos, ectodermal recoil velocity significantly increased in response to NT closure, marking increasing tension in the superficial ectoderm above the NC from non-migratory to early migratory stages (Figure 5a&b), which agrees with recently published data^[63]^. A significant decrease in ectodermal recoil velocity was observed when NT closure was blocked with a Shroom3 morpholino (Figure 5a&b).

**Figure 5:**
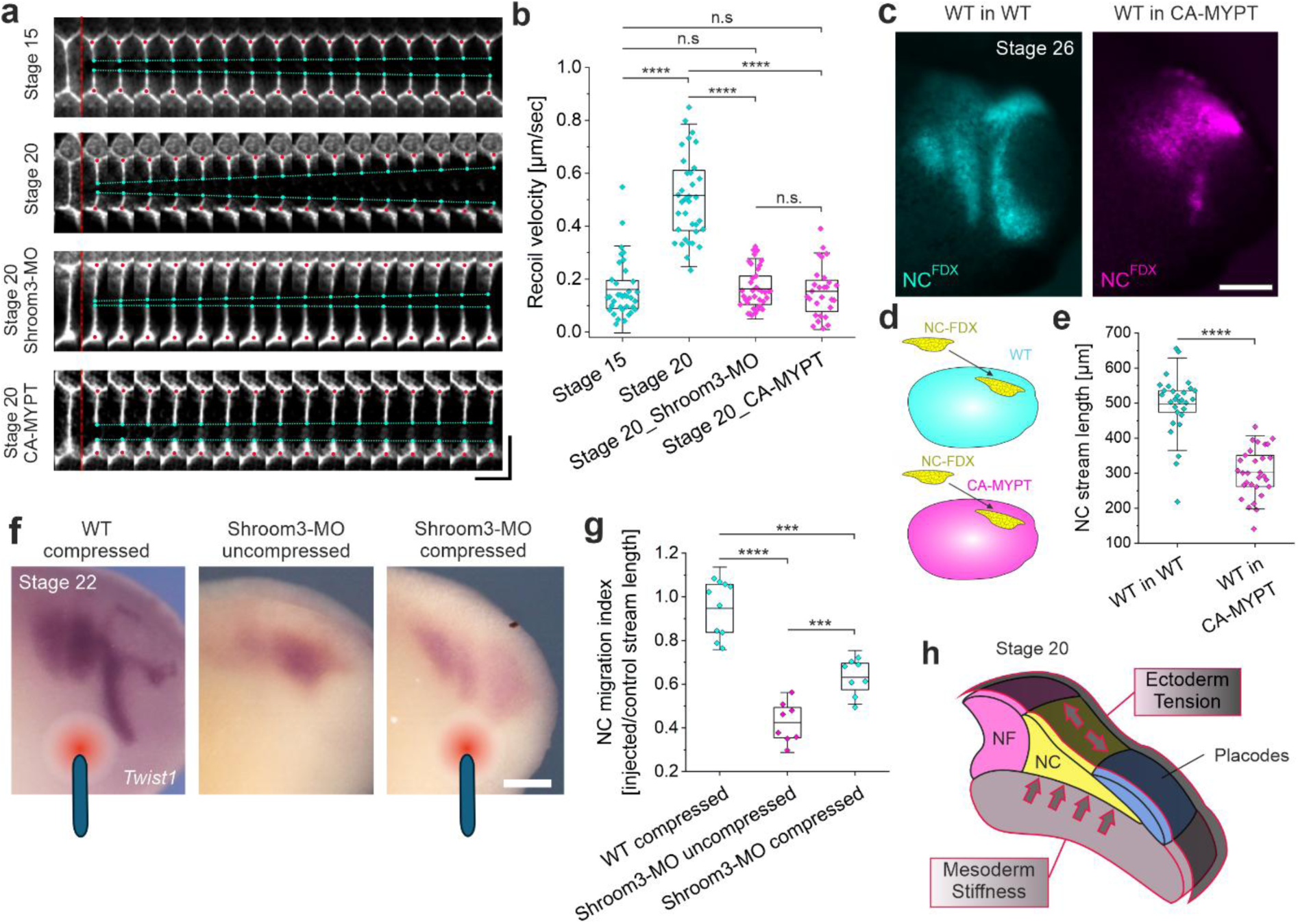
NP bending leads to tension increase in the superficial ectoderm, promoting cephalic NC migration. **a**, Time-lapse montage of cell junction recoil upon laser ablation in Lifeact-RFP expressing embryos before and after NT closure (1^st^ and 2^nd^ row), when NT closure was blocked by Shroom3 morpholino injection (3^rd^ row), or when CA-MYPT was expressed in superficial ectoderm (4^th^ row). Red line marks time of ablation, red dots mark junction vertices, cyan dots mark edges of ablated junction, and dashed cyan lines highlight retraction. Image sequences correspond to Supplement Video 11. **b**, Quantification of ectodermal recoil velocity. **c**, Grafts of fluorescently labeled NC cells into either WT (cyan, left panel) or CA-MYPT (magenta, right panel) hosts. **d**, Schematic illustrating the grafting experiment shown in (**c**). **e**, Length of grafted NC streams at stage 26. **f**, Ectopic tension (marked by blue rods) was exerted on the ventral region ahead of the NC in WT or Shroom3-depleted embryos (left and right panel, compressed) for 7 h. No tension was exerted on Shroom3 morphants as uncompressed control (middle panel). Embryos stained for the NC marker *Twist1* by ISH. **g**, NC migration index was calculated from NC stream length in treated versus contralateral control side. **h**, Schematic illustrating compression of the NC due to mesoderm stiffening and tension increase in the superficial ectoderm, triggered by the bending of the neural fold (NF). Scale bars represent 200 µm (**c**, **f**), and 20 µm (vertical) and 1 sec (horizontal). For **b**, **e**, **g**, data represent mean ± s.d. Statistical analysis was performed using Dunn’s test (**b**), two-tailed *t*-test (**e**), and Tukey’s test (**g**). n.s. *P* > 0.05; \*\*\**P* ≤ 0.001; \*\*\*\**P* ≤ 0.0001. *n* = 36, 25, 28, 28 ablations, respectively (**b**); *n* = 28 control and 30 CA-MYPT grafts (**e**); *n* = 10, 8, 8 embryos, respectively (**g**).

To test if manipulating ectodermal tension reduces NC migration, we expressed constitutively active myosin phosphatase targeting subunit (CA-MYPT) in the prospective placodes and superficial ectoderm by targeted injection into 16-cell blastomeres (Supplement Figure 10f). As expected, we observed a significant reduction in tension in CA-MYPT ectoderm in laser ablation experiments (Figure 5a&b, Supplement Figure 10e, and Supplement Video 11). Next, we grafted control NC into CA-MYPT hosts at stage 16, showing that NC migrated significantly shorter distances compared to NC cells that were grafted into control hosts (Figure 5c-e). Reciprocally, when we ectopically increased tension in the superficial ectoderm of Shroom3 morphants by exerting pulling forces via a micromanipulator set up over 7 hours (Supplement Figure 10g), NC cell migration was partially restored in these migration-deficient embryos (Figure 5f&g).

These data suggest that upon NP bending, the superficial ectoderm is pulled towards the midline, leading to increased ectodermal tension due to stretching reinforcement^[22, 58, 63]^. Together, increasing ectodermal tension and mesodermal stiffness mechanically compress NC cells as they become wedged between the two tissues (Figure 5h).

### Compression primes cephalic NC cells for EMT and FN remodeling

Finally, we set out to understand how compression could tune the migratory behavior of NC cells. NC cells undergo EMT to become motile, and EMT-related genes respond particularly sensitively to mechanical compression in various systems, including invasive cancer cells and *Drosophila*^[39, 64–66]^. Moreover, EMT and ECM remodeling are tightly coupled, with many enzymes linked to ECM remodeling serving as markers for EMT^[67–69]^. Compression arising from NT closure could therefore promote the onset of EMT-related transcriptional programs and mechanosensitive ECM remodeling in NC cells.

To test this hypothesis, we performed RNA-Seq to screen for potential candidate genes that are important for EMT and ECM remodeling, and whose expression is altered upon compression of NC cells. We differentiated NC cells from human induced pluripotent stem cells and performed a transwell migration assay, in which cells squeeze through a porous membrane with pore diameters smaller than a cell’s nucleus. Transwell migration thus requires nuclear deformation leading to nucleus compression (Supplement Figure 11a). Fractions of cells that spontaneously migrated through pores were collected, while the fraction that did not squeeze through pores served as uncompressed control. We observed a total of 268 significantly upregulated genes, among them classic NC EMT-related transcription factors like *Twist1* and *Ets1* (Supplement Figure 11b). Strikingly, among these significantly upregulated genes was *Mmp14* (Supplement Figure 11b), suggesting that onset of NC-mediated ECM remodeling is indeed mechanosensitive.

To further validate this hypothesis, we performed a series of experiments in *Xenopus* embryos. First, we monitored the expression of MMP14 during neurulation in control and Shroom3-depleted embryos. In WT embryos, the onset of MMP14 expression follows shortly after the onset of NP bending and correlates with the onset of NC EMT. From stage 16 onward, MMP14 expression was evident in the cephalic NC and maintained throughout the migratory phase^[36]^ (Supplement Figure 12a). When NT closure was blocked by a Shroom3 morpholino, MMP14 expression was drastically decreased in the cephalic NC (Figure 6a-d). Overexpression of MMP14 in Shroom3 morphants partially rescued their NC migration defect (Supplement Figure 12b-d) and the FN deposition along the NC-NP interface (Figure 6e-h). Strikingly, Shroom3-depleted embryos overexpressing MMP14 showed significantly improved NT morphogenesis, despite continued inhibition of apical constriction (Figure 6e,f,i). These findings further support the idea that FN deposition at the NC-NP interface, dependent on MMP14-NC migration, is required for NT closure.

**Figure 6:**
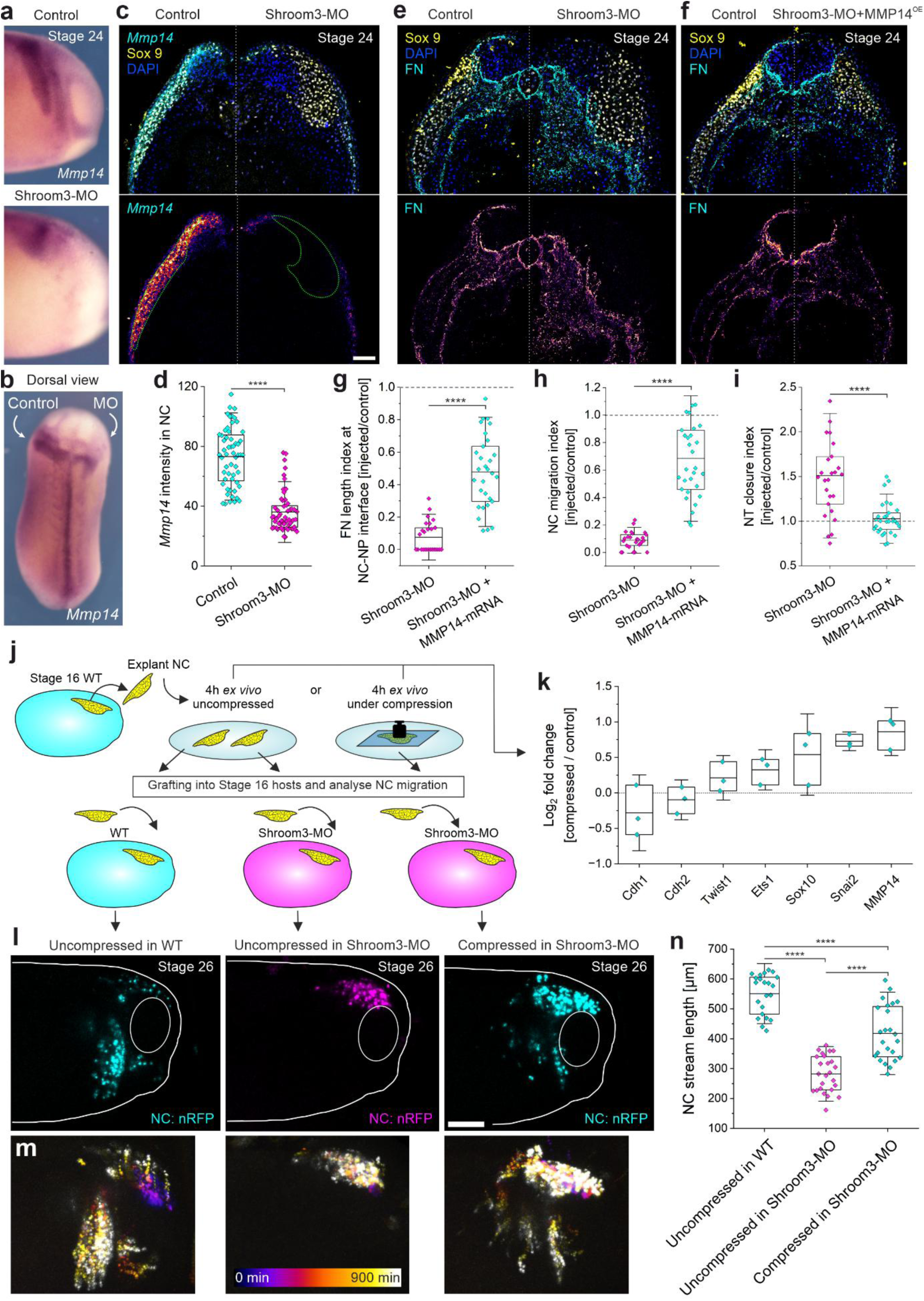
Compression primes cephalic NC cells for FN remodeling by triggering mechanosensitive expression of MMP14 and EMT-related transcription factors. **a**, *Mmp14* expression analyzed at stage 24 by ISH in control and Shroom3-depleted embryo half. **b**, Dorsal view of the embryo depicted in (**a**). **c**, Transverse section of a stage 24 embryo where Shroom3 was depleted in the right side of the embryo. *Mmp14* mRNA was visualized by fluorescent ISH (cyan and intensity LUT in lower panel) and NC were marked by Sox9 (yellow) staining. Nuclei were labeled with DAPI. Green ROI in lower panel marks NC region, defined by Sox9. **d**, *Mmp14* intensity quantification. **e**-**f**, Transverse sections of stage 24 embryos where Shroom3 was depleted from the right side of the embryo (**e**) and where MMP14 was additionally overexpressed (Shroom3-MO + MMP14^OE^) (**f**). Sections were stained for the NC marker Sox9 (yellow) and FN (cyan and intensity LUT in lower panels). **g**-**i**, Quantification of FN length index at NC-NP interface (**g**), NC migration index (**h**), and NT closure index (**i**) in Shroom3-MO or Shroom3-MO + MMP14^OE^ embryos. Indices were calculated by dividing values of injected versus control embryo halves. **j**, Schematic illustrating ex vivo NC compression assay for subsequent qPCR analysis (**k**) or grafting experiments (**l**-**n**). **k**, Differential gene expression of NC EMT-related factors upon compression. Gene expression was measured by qPCR and data are shown as log_2_ fold change of uncompressed versus compressed. **l**, Grafts of fluorescently labeled NC cells at stage 26, corresponding to Supplement Video 12. Uncompressed or compressed NC explants were grafted in either WT or Shroom3-depleted embryos, as illustrated in (**j**). **m**, Temporally color-coded projections of NC streams over 15 h. **n**, Length of grafted NC streams quantified at stage 26. Scale bar in (**c**) represents 100 µm, corresponding to **c**, **e**, **f**; scale bar in (**l**) represents 200 µm. For **d**, **g**, **h**, **i**, **k**, **n**, data represent mean ± s.d. Statistical analysis was performed using two-tailed Mann-Whitney test (**d, g**-**i**) and Tukey’s test (**n**); \*\*\*\**P* ≤ 0.0001. *n* = 50 sections from 10 embryos (**d**); *n* = 26 Shroom3-MO and 30 Shroom3-MO + MMP14^OE^ sections from 11 embryos (**g**-**i**); *n* = 22, 25, 24 embryos, respectively (**n**).

To ensure that these effects are NC cell-autonomous, we grafted MMP14-overexpressing cephalic NC into Shroom3 morphant hosts. Like whole mount overexpression, this was sufficient to restore NC migration in Shroom3-deficient embryos, which don’t undergo NT closure (Supplement Figure 12e&f). Of note, whole mount overexpression of MMP14 in control embryos lead to a slightly premature onset of NC migration (Supplement Figure 12g-i), as reported previously^[36]^, but had little to no effect on NC stream formation, NP bending, or FN remodeling (Supplement Figure 12j-m).

Finally, to test if compression alone is sufficient to increase expression of MMP14 and other EMT-related genes, cephalic NC cells were explanted at the onset of NP bending, compressed ex vivo for 4 hours (Figure 6j), and subsequently analyzed via qPCR. Comparing the expression of EMT-related genes that were upregulated in our RNA-Seq screen revealed increased levels of several candidates upon compression, with *Mmp14* among the strongest responders, together with the transcription factors *Sox10* and *Snai2*, while *Twist1* and *Ets1* showed more moderate responses (Figure 6k). Of note, the classic cadherins, whose switch has been shown to play a prominent role in NC delamination and EMT^[70]^, showed a less pronounced effect in both the human RNA-Seq and *Xenopus* qPCR. While transcripts encoding E-cadherin (*cdh1*) were moderately downregulated in *Xenopus*, which is in line with the idea of a switch from E- to N-cadherin during EMT, transcript levels for N-cadherin (*cdh2*) did not differ between control and confined conditions for both human and *Xenopus* NC cells (Figure 6k and Supplement Figure 11b). This opens the possibility that a switch from E-to N-cadherin is regulated on the post-transcriptional level or that downregulation of E-cadherin is sufficient to shift the balance.

To test if these observed changes in gene expression are sufficient to trigger NC migration in migration-deficient embryos, we repeated our ex vivo compression approach and subsequently grafted compressed or uncompressed NC explants in Shroom3 morphants (Figure 6j, lower panels). Strikingly, compression was sufficient to restore migration in migration-deficient host embryos. While uncompressed grafts migrated in control embryos but not in Shroom3 morphants, compressed grafts migrated directionally over significant distances in Shroom3 morphant hosts (Figure 6l-n and Supplement Video 12), demonstrating that mechanical priming alone is sufficient to induce directed NC migration.

## Discussion

Our findings reveal that cephalic neurulation and NC migration are not sequential morphogenetic events linked by a simple upstream-downstream relationship. Rather, they constitute a reciprocal mechanochemical feedback loop in which each morphogenetic program promotes the progression of the other (Figure 7a). NP bending mechanically compresses cephalic NC cells (Figure 7b), priming them for EMT and MMP14-dependent FN remodeling (Figure 7c). In turn, migrating NC cells remodel FN (Figure 7d) into an extracellular boundary required for continued NT morphogenesis (Figure 7e). Thus, although both NT and NC possess intrinsic genetic programs that drive much of their development autonomously, in vivo the progression of either tissue depends on continuous reciprocal communication with its neighbor.

**Figure 7:**
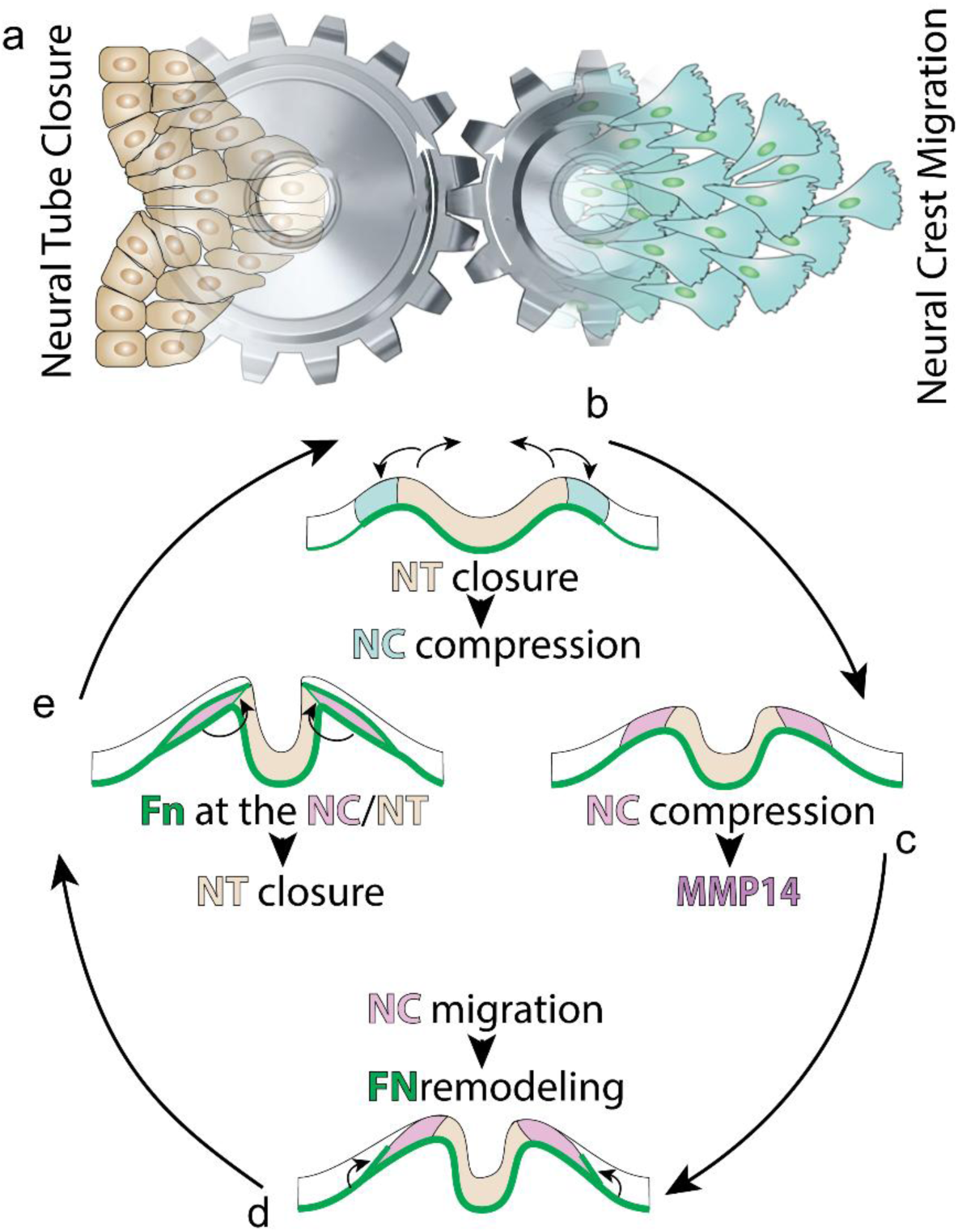
A mechanochemical feedback loop couples NC migration and NT closure in the cephalic region through MMP14-mediated FN remodeling. **a**, Cephalic neurulation and NC migration are not sequential morphogenetic events but constitute a reciprocal feedback loop in which each morphogenetic program promotes the progression of the other. **b**, Bending of the NP leads to mechanical compression of cephalic NC cells. **c**, Compression primes the cephalic NC for EMT and MMP14-dependent FN remodeling. **d**, Migrating NC cells remodel FN into an extracellular boundary along the NC-NP interface and along the NC-ectoderm interface. **e**, FN at the NC-NP interface is not only required for tissue segregation but also for continued NT closure.

A striking feature of this mechanism is that the tissue boundary itself becomes instructive. Rather than acting as a passive partition, the FN-rich NC-NP interface simultaneously maintains tissue segregation and coordinates reciprocal morphogenesis. Unlike mechanisms in which diffusible MMP activity generates broad regions of ECM remodeling^[13, 71, 72]^, MMP14 activity is spatially confined to migrating NC cells. Consequently, ECM remodeling is inseparable from collective cell migration itself, restricting boundary formation to the precise location where tissue separation and morphogenesis occur.

Although this is to our knowledge the first description of this reciprocal feedback mechanism, several observations suggest that its underlying principles may represent conserved developmental modules adapted to species-specific embryonic architecture and timing. NC cells are highly sensitive to tissue mechanics during both delamination^[15, 73]^ and migration^[14, 16]^. Consistent with our finding that compression enhances EMT-related transcriptional programs in *Xenopus laevis*, overcrowding-induced cell extrusion has recently been shown to contribute to NC delamination in mouse embryos through a PIEZO1-dependent mechanism^[73]^. In addition, NC delamination in mouse depends on a mechanical landscape in which cell-shape fluctuations, adhesion, and tissue packing determine the probability of overcoming the energetic barrier to ingression^[74]^. Strikingly, local NC crowding in mouse was linked to NT bending, rapid neuroepithelial proliferation, and embryonic axis elongation^[73]^. Thus, although timing and source of mechanical confinement differ between species, these observations suggest a conserved response of NC cells to compression.

This mechanosensitivity may represent a broader property of highly motile cells, also in the pathological context. Mechanical confinement promotes melanoma invasion^[55]^, while mechanically regulated activation of EMT through *Twist1* has been demonstrated in breast cancer^[39]^. Future work will be required to determine how mechanical compression enhances transcription of *Mmp14* and other EMT-associated genes. Potential mechanisms include changes in chromatin accessibility and nucleocytoplasmic transport, as well as increased nuclear localization of mechanotransducers such as YAP/TAZ, β-catenin, or Twist1 itself^[39, 62, 64, 75]^.

The reciprocal arm of the feedback loop - NC-mediated ECM remodeling as a regulator of NT closure - may likewise represent a conserved developmental module. Although the requirement for NC migration in subsequent NT closure was proposed more than two decades ago^[76, 77]^, its underlying mechanism has remained unclear. *Twist1* is a gene expressed specifically in the NC, and *Twist1*-deficient mice exhibit defects in both cephalic NC migration and NT closure^[78, 79]^, consistent with a functional connection between the two processes. Similarly, recent work in chick suggests that NC-mediated ECM remodeling contributes to cephalic NT expansion^[80]^. The molecular implementation of this feedback may vary according to species-specific tissue architecture while retaining a conserved role for NC-mediated ECM remodeling. For example, chick and mouse embryos establish a collagen-rich basement membrane around the NT earlier than *Xenopus*^[81, 82]^, requiring additional MMPs. Accordingly, chick and mouse cephalic NC additionally express the gelatinases MMP2 and MMP9^[83]^ together with the NC-anchored MMP14^[84]^. However, both MMPs are expressed from the NC rather than the NT. Thus, while the specific proteases involved may differ between species, the underlying principle of NC-dependent ECM remodeling appears to represent a conserved module during neurulation.

FN itself provides a further conserved component of this mechanism. Reduced FN expression causes severe neurulation defects in mice^[74, 85]^, while FN expression or remodeling is required for radial intercalation ^[52]^ and convergent extension^[50, 51, 86]^ in *Xenopus* mesoderm. Moreover, FN can induce tissue segregation in avian ex vivo cultures^[9]^ and promote cell polarization and intercalation in embryonic mouse and zebrafish heart tissue^[53, 54]^. Together, these observations suggest that FN is not merely a structural ECM component but can act as an instructive interface through which tissue boundaries and morphogenetic behaviors are coordinated.

More broadly, our findings have implications for approaches that conceptualize development as the assembly of relatively autonomous tissue modules. The self-organizing capacity of organoids demonstrates the remarkable autonomy of developmental programs^[87]^. Our findings highlight a complementary principle, namely that developmental progression can depend critically on interactions between such programs, with one tissue mechanically and chemically modifying the environment required by its neighbor. Incorporating such reciprocal interactions into organoid and tissue-engineering systems may therefore be essential for reproducing morphogenesis that depends not only on intrinsic tissue programs, but also on dynamic communication between adjacent developing tissues.

## Supplementary Figures

**Supplement Figure 1:**
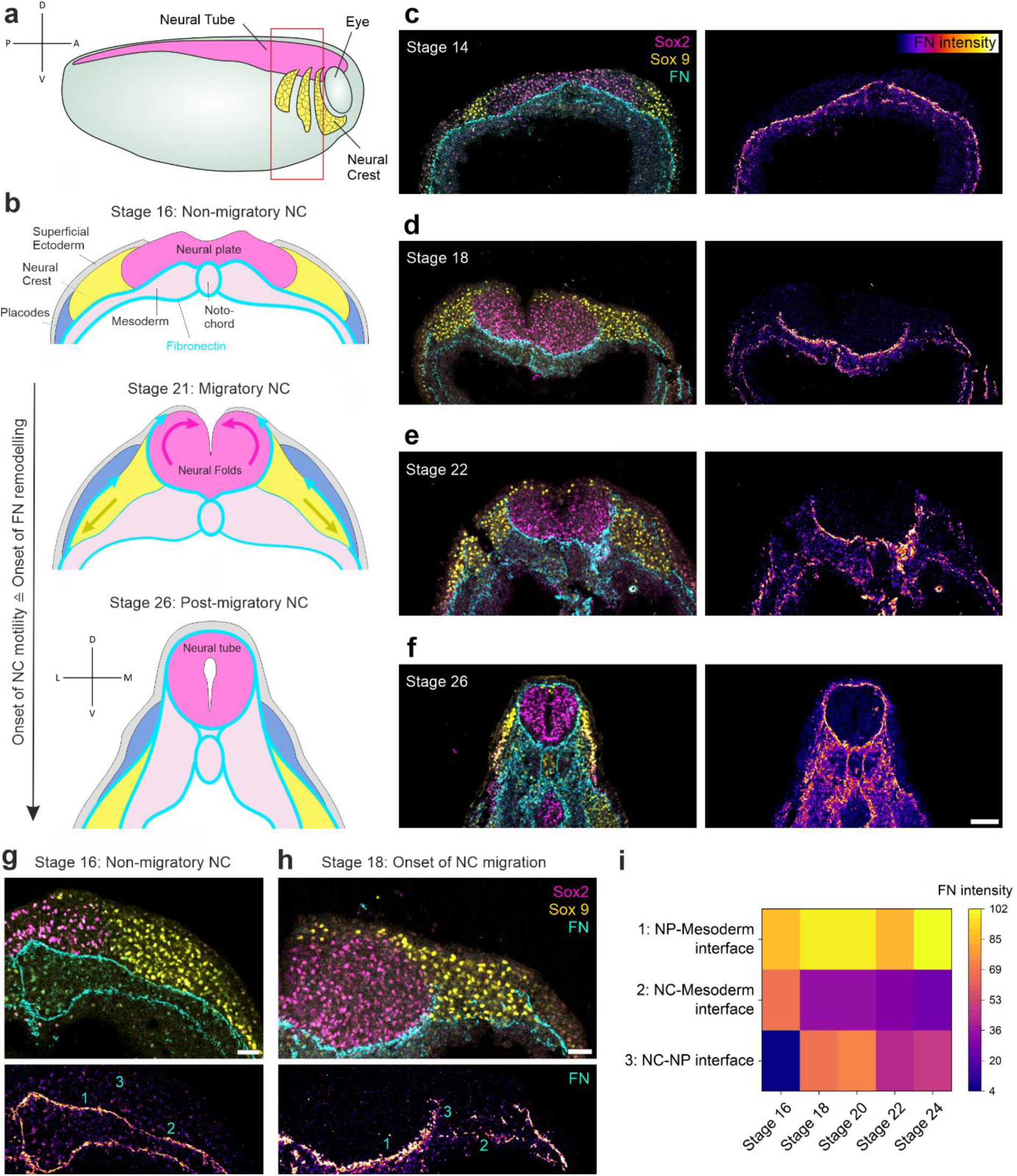
Dynamics of FN remodeling during cephalic NT closure and NC migration. **a**, Lateral view schematic of embryo with cephalic NC streams (yellow) and NT (magenta). Red box marking region from which cryosections were analyzed. **b**, Schematic illustration of FN remodeling during cephalic NT closure and NC migration (transverse view), corresponding to ROI highlighted by the red box in (**a**). **c**-**f**, Transverse sections of stage 14 (**c**), 18 (**d**), 22 (**e**), and 26 (**f**) embryos, stained for the NP marker Sox2 (magenta), the NC marker Sox9 (yellow), and FN (cyan and LUT in the right panels). **g**-**h**, Close-up views from transverse sections, showing NP (Sox2, magenta), NC (Sox9, yellow), and FN (cyan and LUT in lower panels) before (**g**) and after onset of NC migration (**h**). Numbers indicate ROIs for FN intensity measurements along the NP-mesoderm interface (1), NC-mesoderm interface (2), and NC-NP interface (3). **i**, Heatmap of mean FN intensities along the indicated interfaces over developmental stages 16-24. *n* = 32-45 measurements from 10 different embryos (**i**). Scale bar in (**f**) represents 100 µm (for **c**-**f**), and 50 µm for (**g**) and (**h**).

**Supplement Figure 2:**
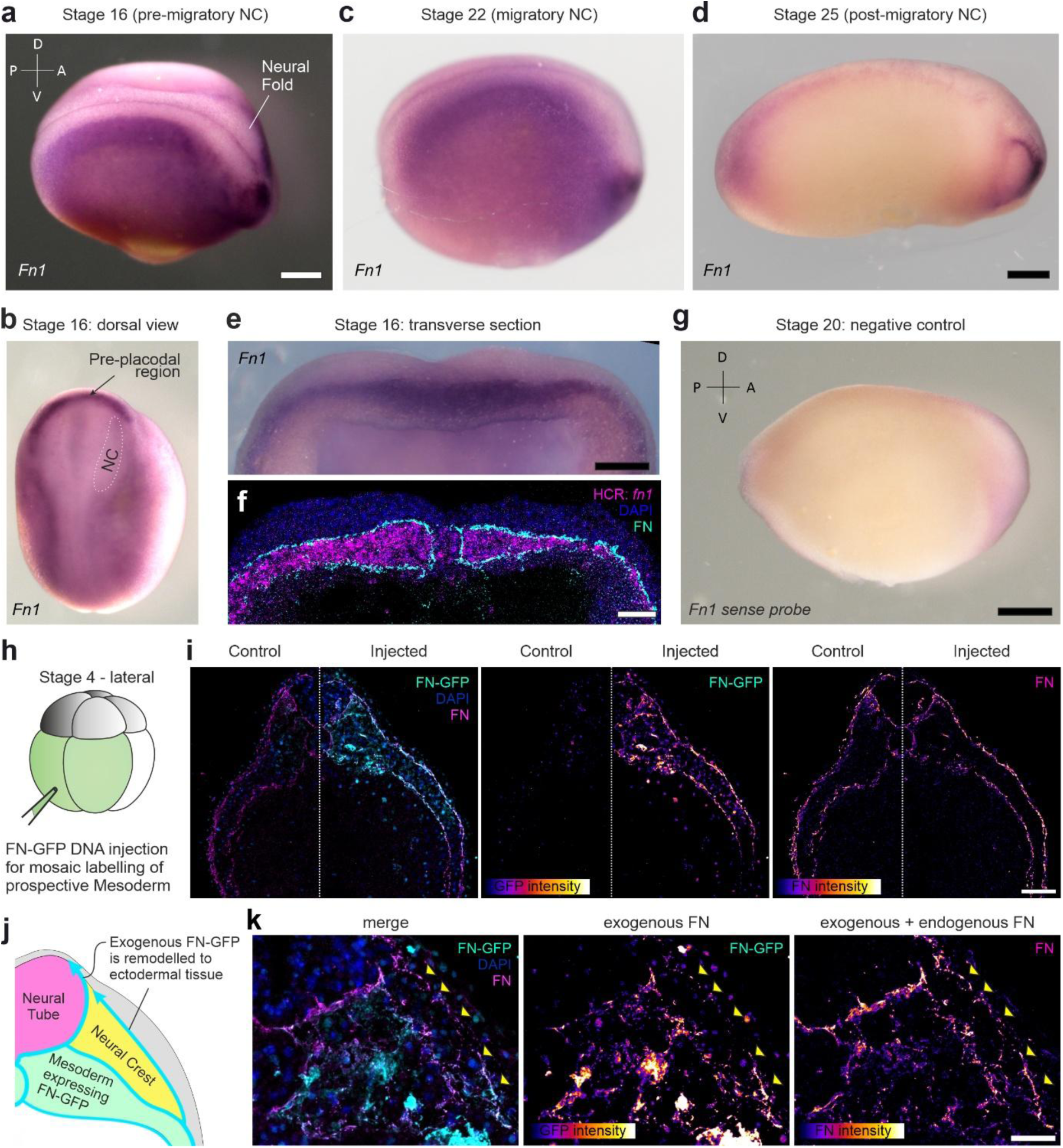
Mesoderm-derived *Fn1* expression and FN protein distribution during NC migration. **a**-**d**, Colorimetric ISH against *Fn1* at pre-migratory (**a**&**b**), migratory (**c**), and post-migratory NC stages (**d**). **b**, Dorsal view of embryo shown in (**a**). Crescent-shaped staining at anterior end corresponds to pre-placodal region. **e**, Transverse section of stage 16 embryo, stained for *Fn1* by colorimetric ISH. **f**, Transverse section of stage 16 embryo, stained for *Fn1* by fluorescent HCR (magenta), with subsequent staining of FN protein (cyan) and nuclei (DAPI). **g**, Mock control (sense probe) for colorimetric ISH staining of *Fn1*. **h**, Schematic illustrating injection of DNA encoding FN-GFP at Stage 4 into the ventral blastomeres, giving rise to mesoderm. **i**, Transverse section of stage 24 embryo with FN-GFP expression in right embryo half, showing that exogenous FN-GFP reconstitutes localization of endogenous FN. Left panel shows merge image with endogenous FN in magenta and FN-GFP in cyan. Nuclei were additionally stained with DAPI. Middle panel shows FN-GFP intensity, right panel endogenous FN intensity as LUT. **j**, Schematic highlighting redistribution of exogenous mesoderm-derived FN along NC-ectoderm and NC-NP interfaces. **k**, Redistribution of exogenous mesoderm-derived FN along NC-ectoderm interface in vivo. Arrows mark FN-GFP at the NC-ectoderm interface and the NC area underneath. Left panel shows merged image with endogenous FN in magenta and FN-GFP in cyan. Middle panel shows the FN-GFP intensity and right panel endogenous FN intensity as LUT. GFP in (**i** and **k**) was additionally stained with an Alexa Fluor 488-coupled antibody to enhance contrast. Scale bars represent 200 µm in **a**, **b**, **c**, **d**, **e**, **g**; 100 µm in **f**, **i**; 50 µm in **k**.

**Supplement Figure 3:**
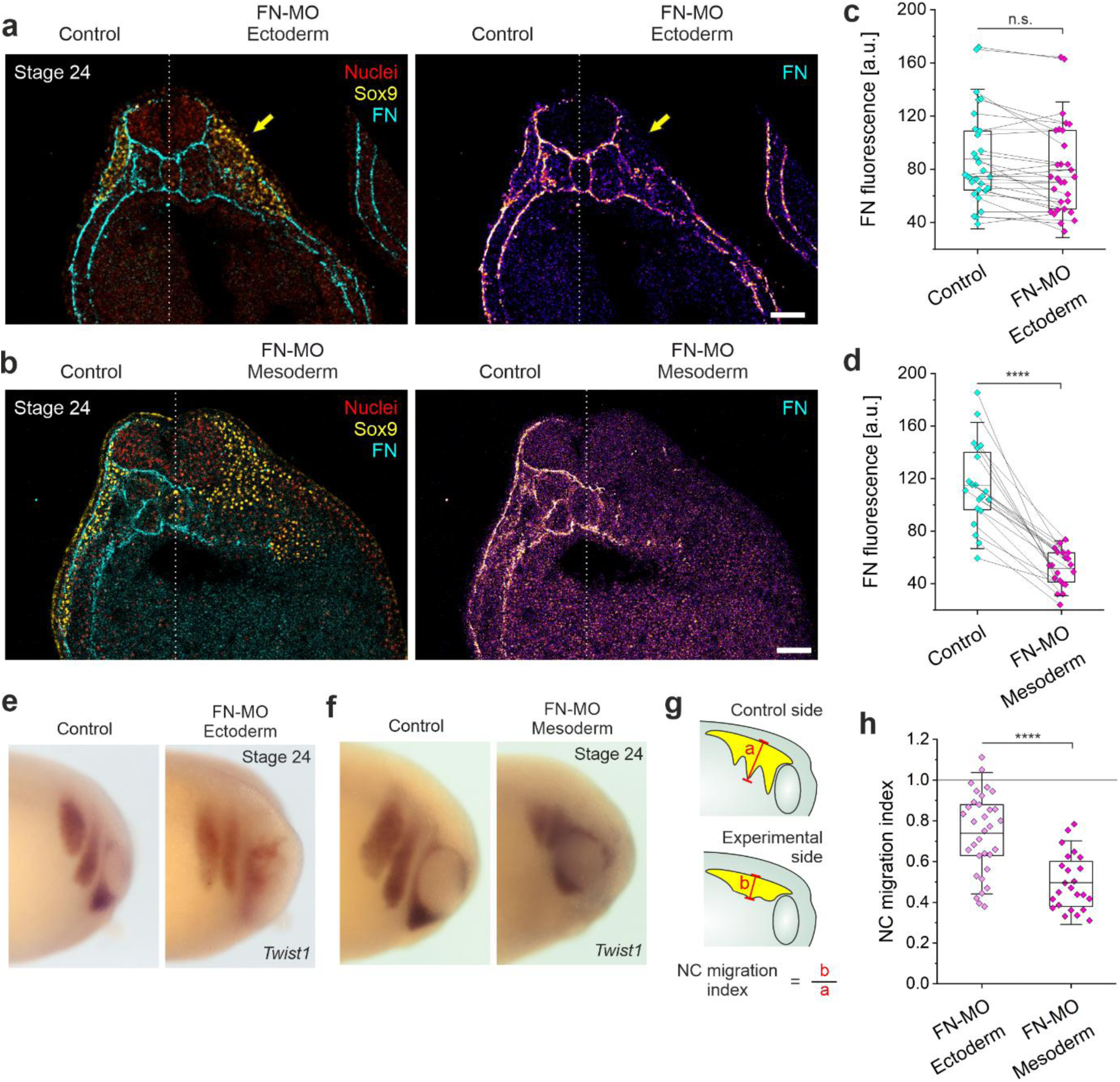
Effect of FN depletion from the mesoderm or placodes on FN distribution and NC migration. **a**-**b**, Transverse section of stage 24 embryo, in which FN expression from placodes (**a**) or mesoderm (**b**) was blocked in right side of the embryo. Staining against NC marker Sox9 (yellow), FN (cyan and LUT in right panel). Nuclei (DAPI) are shown in red. **a**, Depletion of FN from placodes has little effect and mainly leads to shorter FN at the NC-ectoderm interface (yellow arrow), while FN at the NC-NP interface remains unaffected. **b**, Depletion of FN from the mesoderm leads to almost complete absence of FN from the right side of the embryo. **c**&**d**, Mean FN intensity, when FN was depleted from placodes (**c**) or the mesoderm (**d**). **e**-**f**, Embryos were injected with *Fn1* morpholino in one side, depleting FN from the placodes (**e**) or mesoderm (**f**), and stained for the NC marker *Twist1* by ISH. **g**, Schematic illustrating quantification of NC stream length and calculation of NC migration index. **h**, NC migration index for embryos where FN was depleted from placodes or the mesoderm. Scale bar represents 100 µm in (**a**) and (**b**). For **c**, **d**, **h**, data are mean ± s.d. Statistical analysis was performed using two-tailed Mann-Whitney test (**c**) and two-tailed *t*-test (**d**, **h**); n.s. *P* > 0.05; \*\*\*\**P* ≤ 0.0001. *n* = 30 measurements from 10 different embryos (**c**); *n* = 20 measurements from 10 different embryos (**d**); *n* = 33 (FN-MO Ectoderm) and 24 (FN-MO Mesoderm) embryos (**h**).

**Supplement Figure 4:**
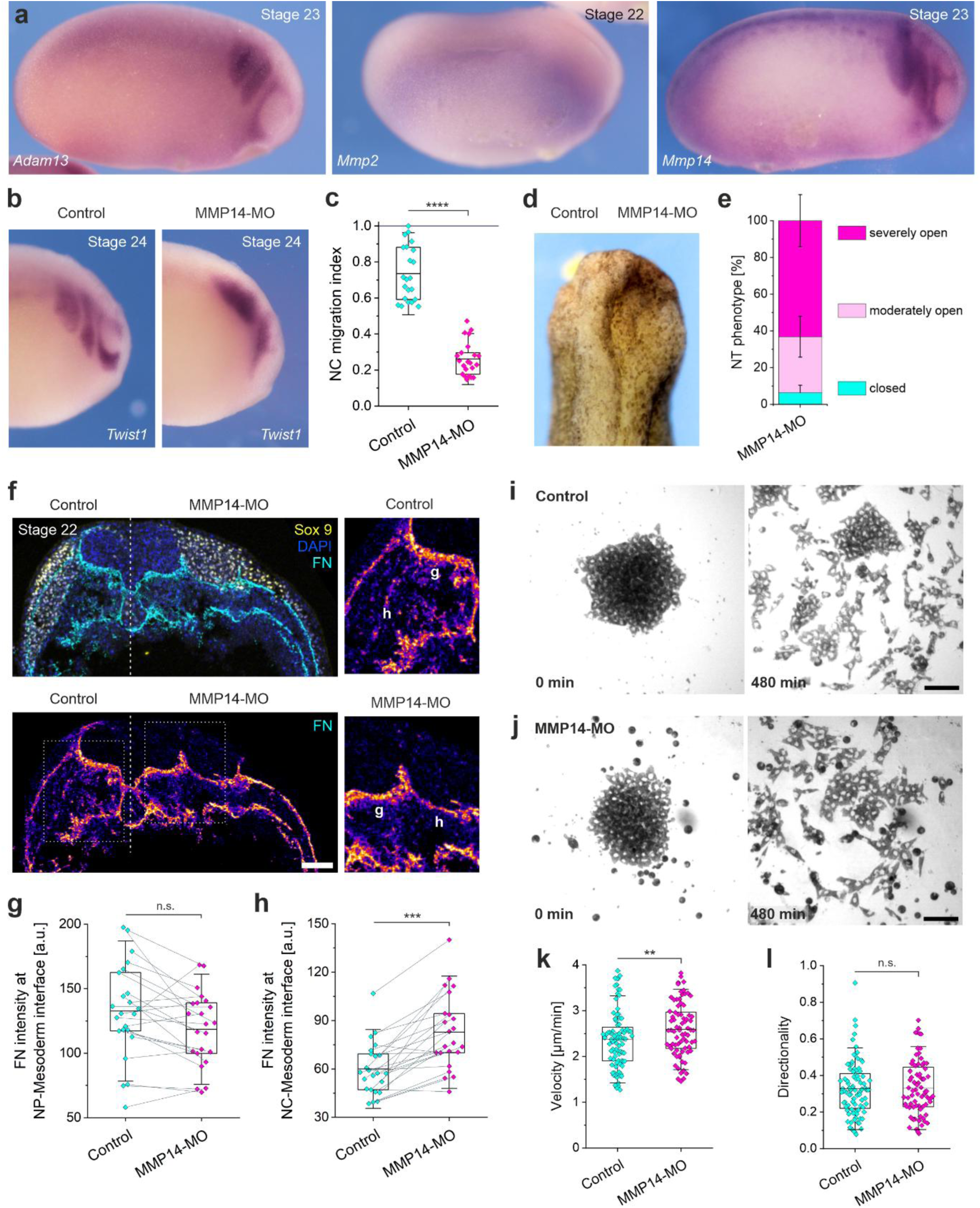
MMP14 is necessary for FN remodeling during NC invasion in vivo. **a**, ISH against *Adam 13* (left panel), *Mmp2* (middle panel), and *Mmp14* (right panel) during migratory NC stages. **b**, Embryos were injected with *Mmp14* morpholino in one side and stained for the NC marker *Twist1* by ISH. **c**, NC migration index in MMP14-depleted embryos. NC stream length were normalized to the maximum value in the control. **d**, Dorsal view of stage 24 embryo, in which MMP14 was depleted from right side of the embryo, leaving the NT open. **e**, Ratio of phenotypes, categorized by visual inspection of cephalic NT, upon depletion of MMP14 from one embryo half. **f**, Transverse section of stage 22 embryo, in which MMP14 expression was blocked in right side of the embryo. Staining against Sox9 (yellow), FN (cyan and LUT in lower panel) and nuclei (DAPI). Right panels showing FN intensity as LUT correspond to the boxes highlighted in the lower panel of the overview. Labels highlight the FN interfaces from which measurements were taken. **g**-**h**, Mean FN intensities measured along the NP-mesoderm (**g**) and NC-mesoderm (**h**) interface in control and MMP14-depleted embryo halves. **i**-**j**, Still images of control (**i**) and MMP14-depleted (**j**) NC explants, migrating on FN-coated dishes. Images show NC cell dispersion at 0 min (left panels) and after 480 min (right panels), corresponding to Supplement Video 1. **k**-**l**, Velocity (**k**) and Directionality (**l**) of single control or MMP14-depleted NC cells. Part of control dispersion data in (**k**) and (**l**) are re-plotted in Supplement Figure 9c-f. Scale bars represent 100 µm in **f**, **i**, **j**. For **c**, **e**, **g**, **h**, **k**, **l**, data are mean ± s.d. Statistical analysis was performed using two-tailed *t*-test (**c**, **g**, **h**) and two-tailed Mann-Whitney test (**k**, **l**); n.s. *P* > 0.05; \*\**P* ≤ 0.01; \*\*\**P* ≤ 0.001; \*\*\*\**P* ≤ 0.0001. *n* = 22 embryos (**c**); *n* = 95 embryos (**e**); *n* = 23 measurements from 12 different embryos (**g**, **h**); *n* = 88 control and 76 MMP14-depleted cells (**k**, **l**).

**Supplement Figure 5:**
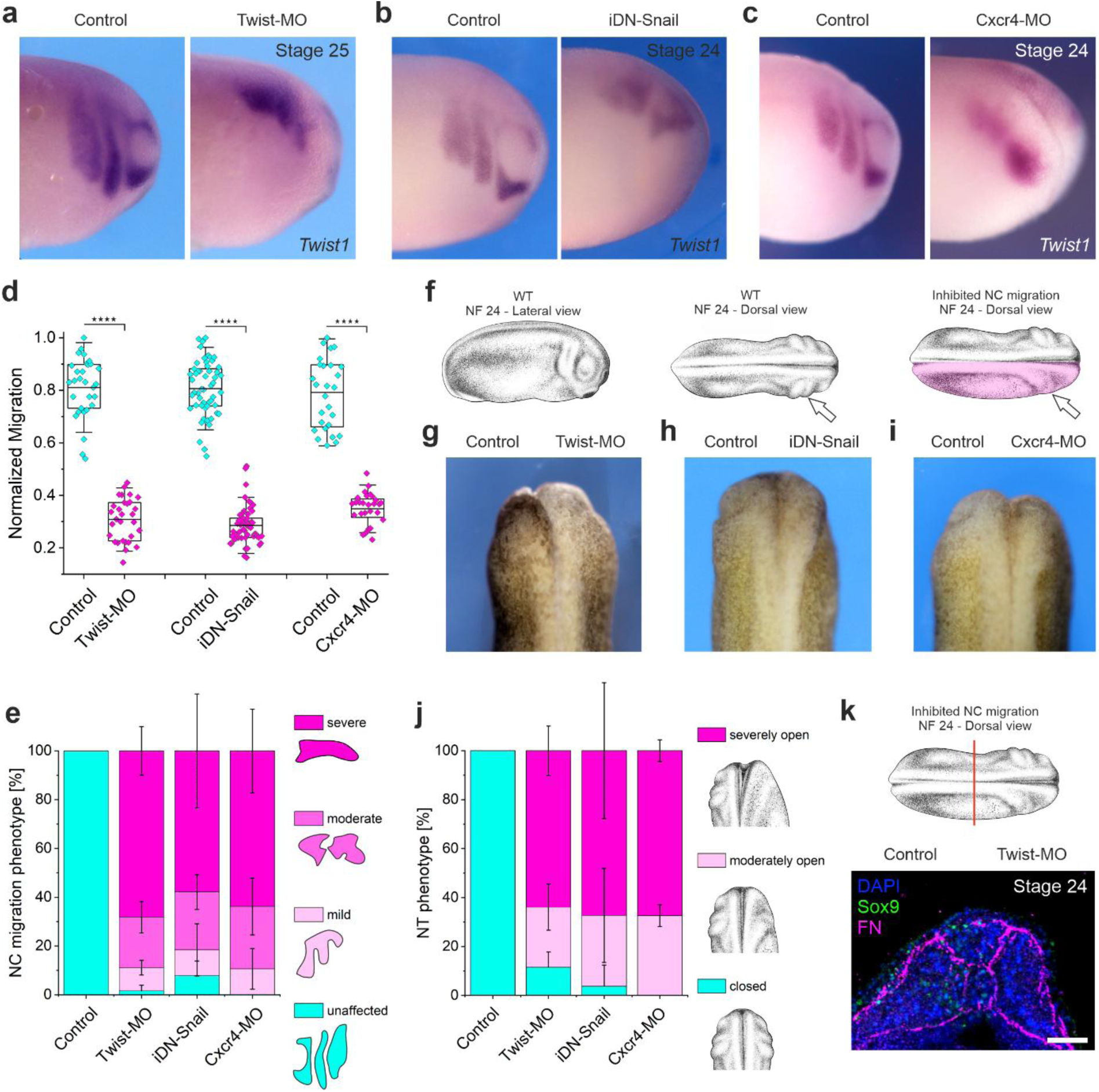
Blocking NC migration simultaneously inhibits NT closure. **a**-**c**, Embryos were injected into one side with a *Twist1* morpholino (**a**), inducible dominant negative Snail (**b**), or a *Cxcr4* morpholino (**c**), and the NC marker *Twist1* was stained by ISH. **d**, NC migration index in the respective embryos. NC stream length were normalized to the maximum value in the control. **e**, Ratios of NC migration phenotypes, categorized by criteria depicted in the legend. **f**, Illustrations depicting development of the branchial arches (arrow) upon NC migration and NT closure. Note that upon inhibition of NC migration, branchial arches do not become visible as striking features (arrow in right panel). Xenopus illustrations © Natalya Zahn (2022)^[89]^. **g**-**i**, Dorsal view of stage 24 embryos, in which NC migration was blocked in the right side through injection of *Twist1* morpholino (**g**), inducible dominant negative Snail (**h**), or *Cxcr4* morpholino (**i**), leaving the NT open. **j**, Ratios of NT phenotypes, categorized by the criteria depicted in the legend. **k**, Transverse section through posterior region of a stage 24 embryo, as depicted in schematic. Cephalic NC migration was blocked by injecting *Twist1* morpholino into the right side of the embryo. Section was stained for Sox9 (green), FN (magenta), and nuclei (DAPI). Xenopus illustration © Natalya Zahn (2022)^[89]^. Scale bar in (**k**) represents 100 µm. For **d**, **e**, **j**, data are mean ± s.d. Statistical analysis was performed using two-tailed Mann-Whitney test (**d**); \*\*\*\**P* ≤ 0.0001. *n* = 30 Twist-MO embryos, 54 iDN Snail embryos, and 28 Cxcr4-MO embryos (**d**); *n* = 63 Twist-MO embryos, 76 iDN Snail embryos, and 47 Cxcr4-MO embryos (**e**); *n* = 122 Twist-MO embryos, 107 iDN Snail embryos, and 100 Cxcr4-MO embryos (**j**).

**Supplement Figure 6:**
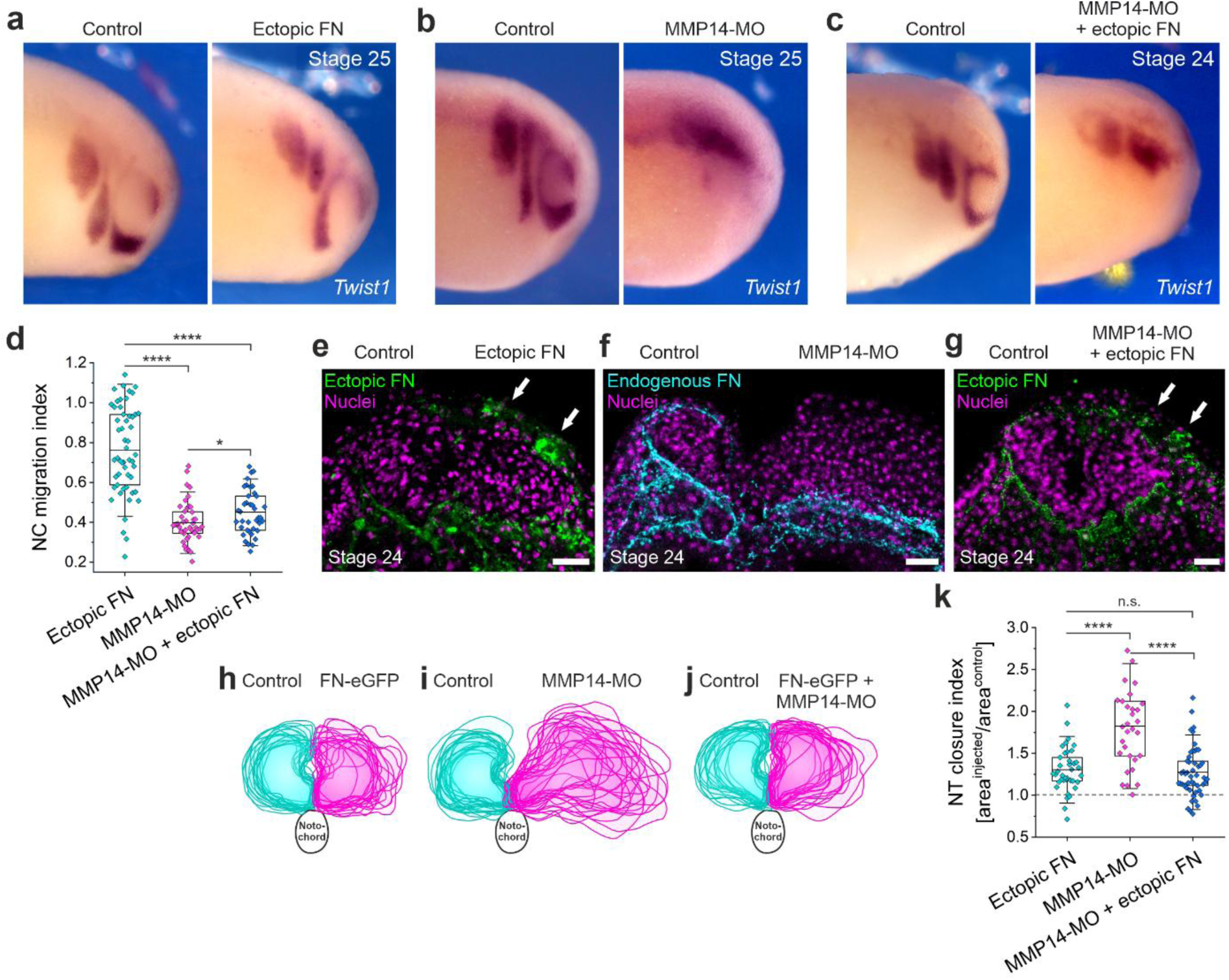
Ectopic FN expression rescues NT morphogenesis in the absence of NC-mediated FN remodeling. **a**-**c**, Stage 24-25 embryos were stained for the NC marker *Twist1* by ISH. **a**, Injection of DNA encoding FN-GFP at stage 5 to express FN ectopically from the superficial ectoderm and placodes; **b**, Injection of *Mmp14* morpholino at stage 4 to block expression from one side of the embryo; **c**, *Mmp14* morpholino injection at stage 4 and subsequent injection of FN-GFP DNA at stage 5 to achieve both effects simultaneously. **d**, NC migration index. NC stream length in the injected embryo half was compared to the respective contralateral control, each. **e**-**g**, Transverse sections of embryos, treated as described for (**a**-**c**). Sections were either stained against GFP (green) to enhance visibility of ectopic FN (**e** and **g**), or against endogenous FN (cyan) (**f**) in MMP14 morphants. Nuclei were stained with DAPI. **h**-**j**, Overlayed areal projections of control (cyan) and injected (magenta) NT hemispheres from 18 FN-eGFP, 21 MMP14-MO, and 18 FN-eGFP + MMP14-MO sections of 10 embryos each. **k**, NT closure index for the respective indicated treatments. Note that an index of 1 indicates similar NT areas (closed NT). Scale bars represent 100 µm for (**e**-**g**). For **d** and **k**, data are mean ± s.d. Statistical analysis was performed using Tukey’s test (**d** and **k**); n.s. *P* > 0.05; \**P* ≤ 0.05; \*\*\*\**P* ≤ 0.0001. *n* = 53 FN-eGFP embryos, 44 MMP14-MO embryos, and 39 FN-eGFP + MMP14-MO embryos (**d**); *n* = 38 FN-eGFP sections, 31 MMP14-MO sections, and 49 FN-eGFP + MMP14-MO sections from 10 different embryos each (**k**).

**Supplement Figure 7:**
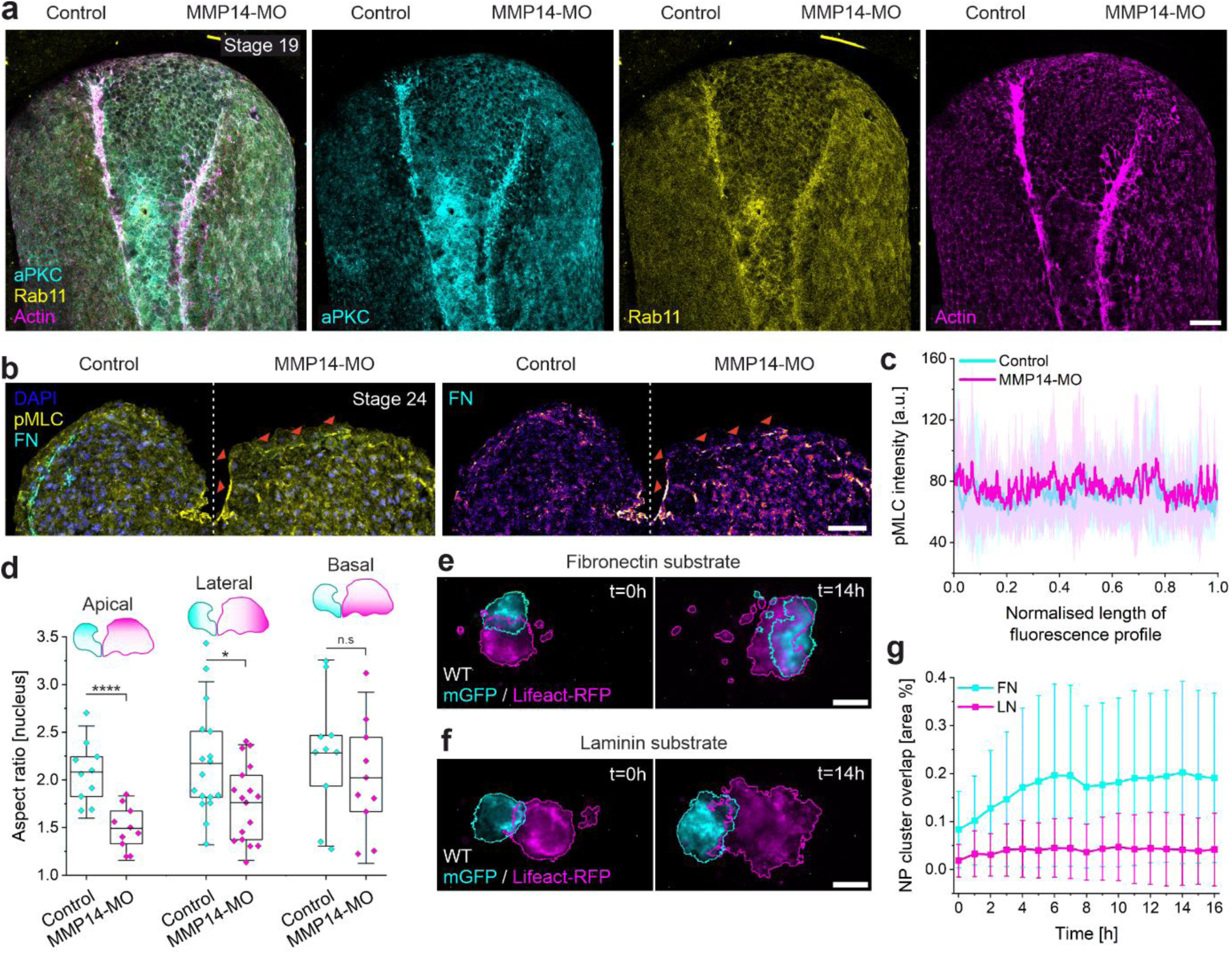
Effects of FN on apical constriction and cell intercalation. **a**, Cephalic region of an embryos at stage 19, in which MMP14-dependent FN remodeling was blocked in the right side of the embryo. The embryo was stained for aPKC (cyan), Rab11 (yellow), and actin (phalloidin, magenta). Left panel shows the merged image, followed by individual channels each. **b**, Transverse section of stage 24 embryo with MMP14 depleted from right side, stained for phosphorylated myosin light chain (pMLC, yellow) and FN (cyan and intensity LUT in right panel). Nuclei were stained with DAPI. Arrowheads indicate pMLC foci along apical surface of the NT. **c**, Intensity profiles of pMLC along apical surface of control and MMP14-depleted NT hemispheres. Length of individual line scans were normalized and line profiles averaged to generate mean ± s.d. profiles. **d**, Aspect ratio of NT nuclei cross-sections in ROIs depicted in legend (apical, lateral, basal) from control and MMP14-depleted embryo half. **e**-**f**, NP cell intercalation on FN (**e**) or LN (**f**) ex vivo. NP explants expressing mGFP or Lifeact-RFP were placed in direct proximity, and intercalation was monitored over time. Still images at t = 0 h (left panels) and t = 14 h (right panels) correspond to Supplement Video 9. **g**, Intercalation of NP explants on FN or, plotted as overlap between mGFP and Lifeact-RFP LN over time. Scale bars represent 100 µm (**a**, **e**, **f**) and 50 µm (**b**). For **c**, **d**, **g**, data are mean ± s.d. Statistical analysis was performed using two-tailed *t*-test (**d**); n.s. *P* > 0.05; \**P* ≤ 0.05; \*\*\*\**P* ≤ 0.0001. *n* = 19 control line scans and 17 MMP14-MO line scans from 10 different embryos (**c**); *n* = 10 apical, 17 lateral, and 10 basal cells from single SEM slice (**d**); *n* = 14 explant pairs on FN and 24 explant pairs on LN (**g**).

**Supplement Figure 8:**
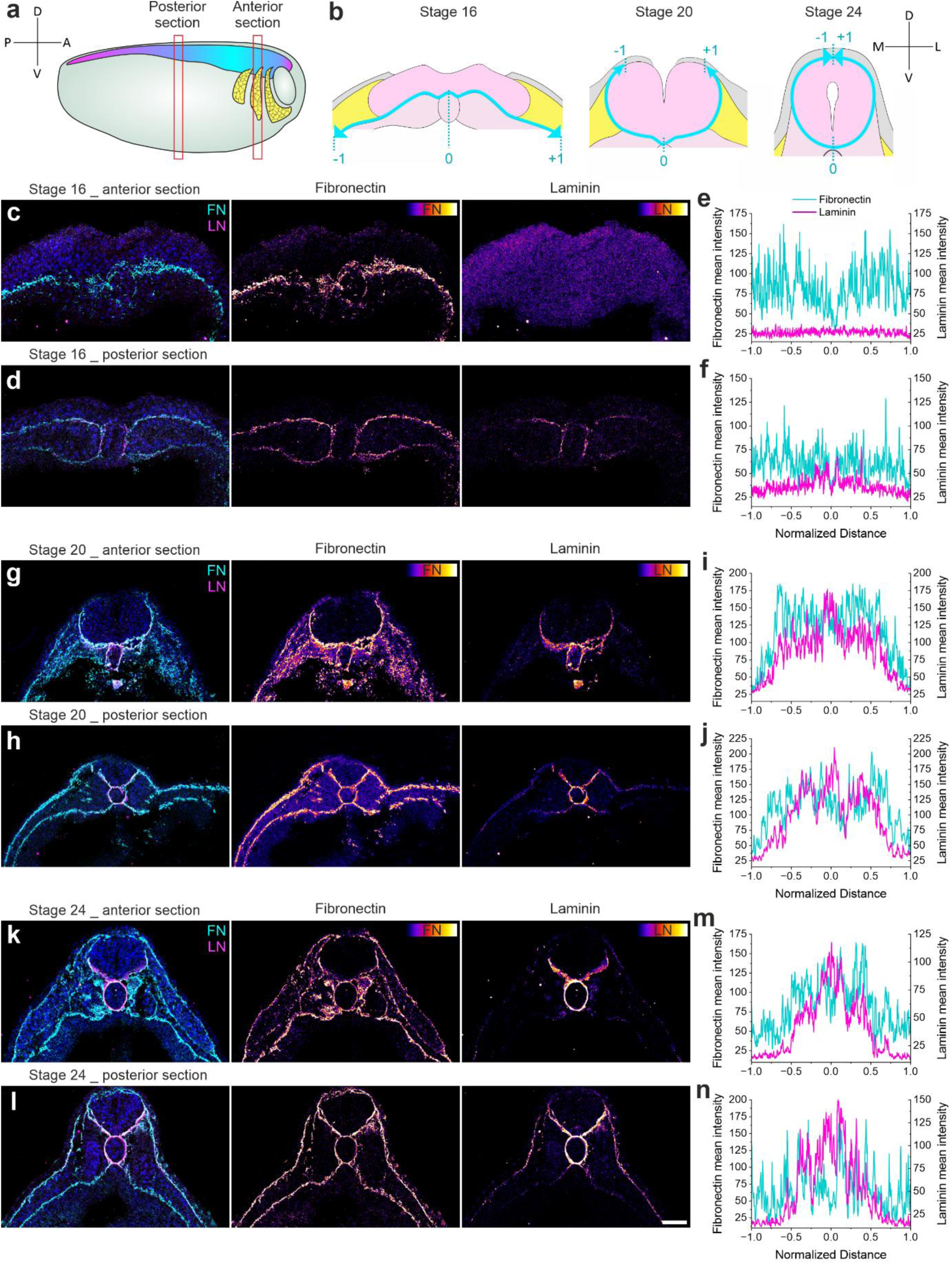
FN and LN localization during cephalic NT closure and NC migration. **a**, Lateral view schematic of an embryo with the cephalic NC streams (yellow) and the NT (magenta/cyan gradient). Red boxes mark ROIs from which cryosections were analyzed. **b**, Transverse view schematic illustrating where line scans were taken at distinct stages. **c**-**l**, Transverse sections through the anterior (**c**, **g**, **k**) and posterior (**d**, **h**, **l**) region at stage 16 (**c** & **d**), stage 20 (**g** & **h**), and stage 24 (**k** & **l**), stained for FN (cyan and intensity LUT in middle panel) and LN (magenta and intensity LUT in right panel). All line profiles shown in (**e** & **f**), (**i** & **j**), (**m** & **n**) correspond to the according stages and regions depicted, and are averaged from multiple sections and embryos, with the length profile being normalized as illustrated in (**b**). Scale bar represents 100 µm for (**c** & **d**), (**g** & **h**), (**k** & **l**). *n* = 5 (**c**), 5 (**d**), 7 (**g**), 5 (**h**), 8 (**k**), and 4 (**l**) sections from 3 embryos each.

**Supplement Figure 9:**
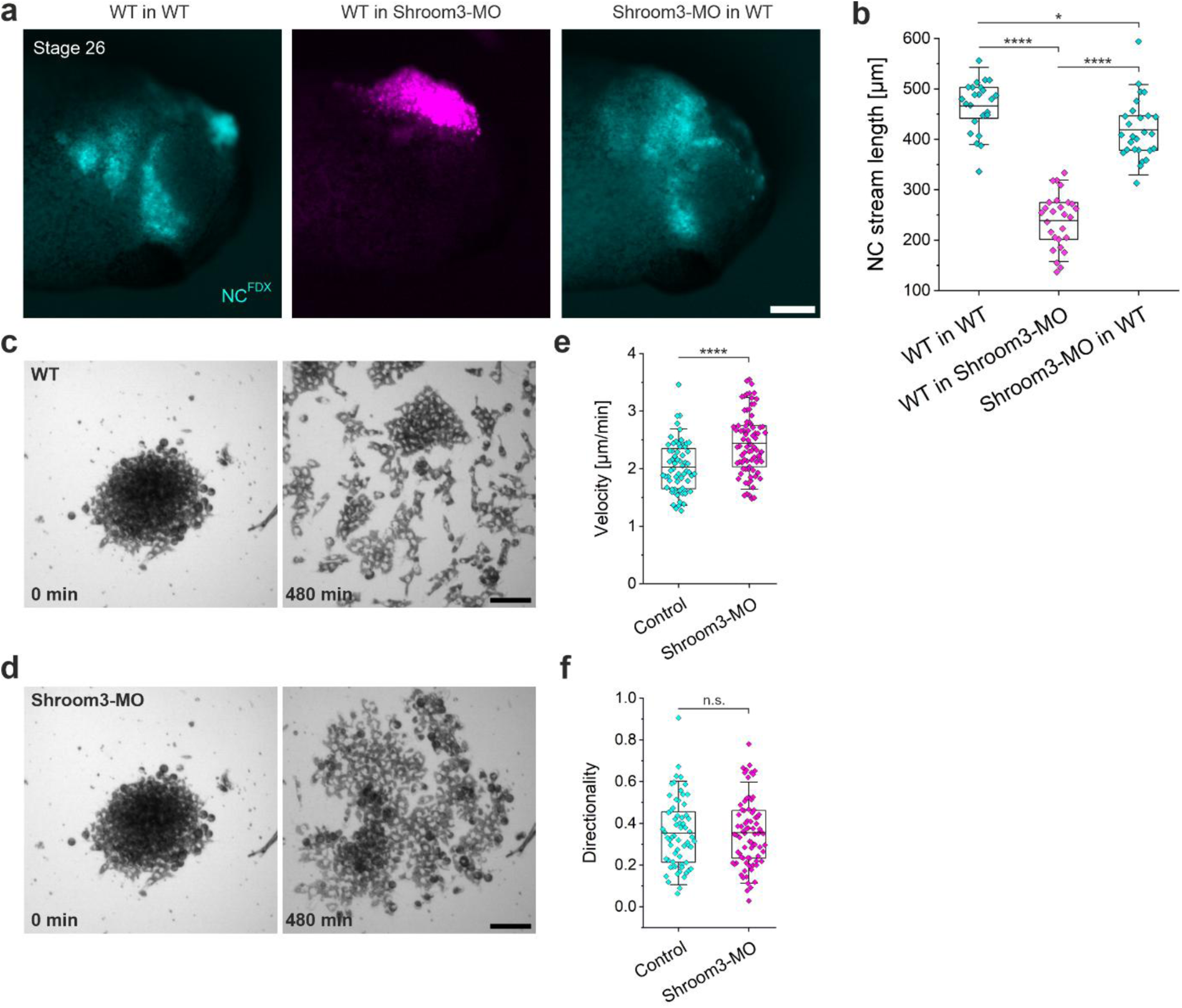
Shroom3 depletion affects NC cells in a non-cell autonomous manner. **a**, Grafts of fluorescently labeled NC cells. WT NC were transplanted either in WT (cyan) or Shroom3 morphant (magenta) hosts (left and middle panel), and NC cells from Shroom3 morphant embryos were transplanted into WT (cyan) hosts (right panel). **b**, Length of grafted NC streams at stage 26. Parts of data for the WT NC grafts in Shroom3-MO hosts are re-plotted in Supplement Figure 12f. **c**-**d**, Still images of NC explants from control (**c**) and Shroom3-depleted (**d**) embryos, migrating on FN-coated dishes. Images show NC cell dispersion at 0 min (left panels) and 480 min (right panels), corresponding to Supplement Video 10. **e**-**f**, Velocity (**e**) and Directionality (**f**) of single NC cells from control or Shroom3-depleted embryos. Part of control dispersion data in (**e**) and (**f**) are re-plotted in Supplement Figure 4k-l. Scale bars represent 200 µm (**a**) and 100 µm (**c** and **d**). For **b**, **e**, **f**, data are mean ± s.d. Statistical analysis was performed using Tukey’s test (**b**) and two-tailed *t*-test (**e**, **f**); n.s. *P* > 0.05; \**P* ≤ 0.05; \*\*\*\**P* ≤ 0.0001. *n* = 24, 27, and 27 embryos, respectively (**b**); *n* = 64 cells from control and 81 cells from Shroom3-depleted embryos (**e**, **f**).

**Supplement Figure 10:**
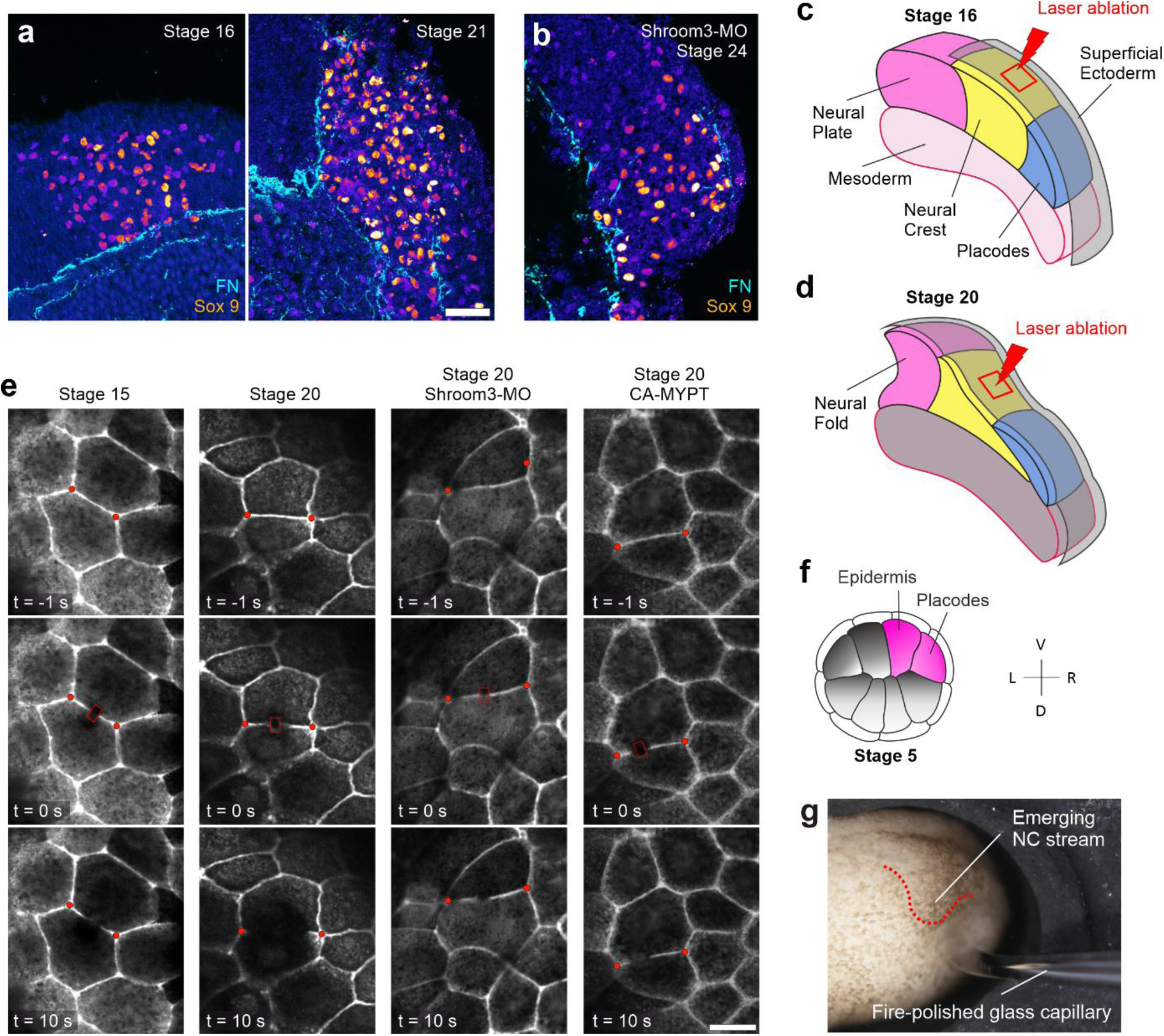
NC compression arises from tension increase in the superficial ectoderm. **a**-**b**, Fields of view, corresponding to NC stream close-up views in Figure 4f. Sections were stained for the NC marker Sox9 (intensity LUT) and FN (cyan). Shroom3 was depleted from the embryo in (**b**). **c**-**d**, Schematics illustrating ROIs for laser ablation. **e**, Still images of cell junction recoil upon laser ablation in Lifeact-RFP expressing embryos, before (top row), at t = 0 sec (middle row), and at t = 10 sec (bottom row) after laser ablation. Red box marks ablation ROI, red dots mark junction vertices. Image sequences correspond to Supplement Video 11. **f**, Schematic illustrating injection of CA-MYPT mCherry at Stage 5 into the animal blastomeres, giving rise to the placodes and superficial ectoderm. **g**, Example image of the compression assay, shown in Figure 5f&g. The glass capillary is held in place by a micromanipulator.

**Supplement Figure 11:**
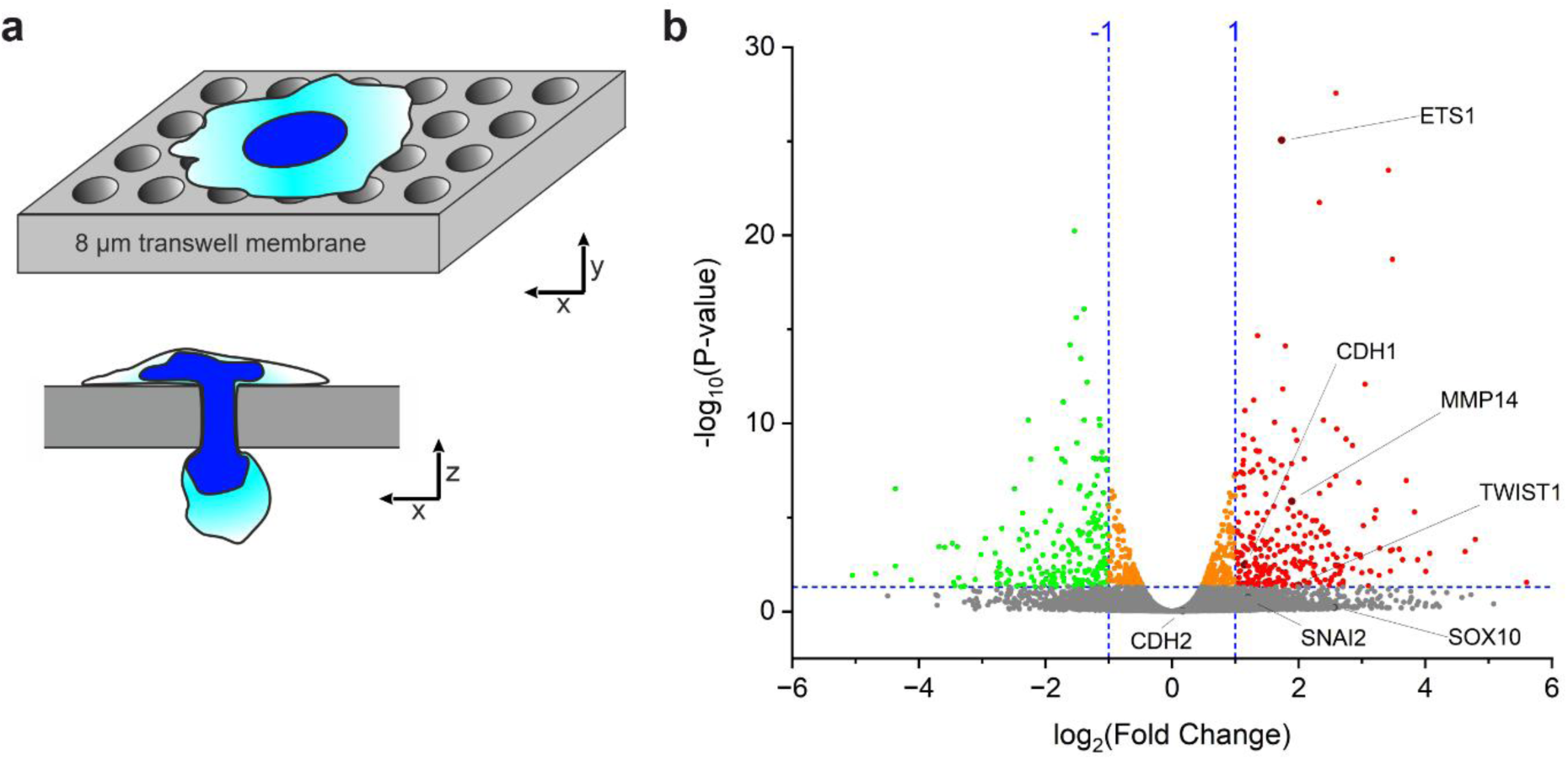
Transcriptional changes in human induced neural crest cells associated with mechanical compression. **a**, Schematic of the transwell assay. To squeeze through the pores, neural crest cells must deform their nucleus, exerting compressive forces onto the nucleus. Cells that migrated through the porous membrane where harvested as compressed fraction, while cells remaining on the upper compartment were collected as uncompressed control fraction. b, Volcano plot showing differential gene expression between uncompressed and compressed fractions. Each dot represents one gene, genes significantly upregulated are shown in red, significantly downregulated genes are shown in green, and genes that show no significant difference are shown in gray. Dashed lines indicate the thresholds for fold change and statistical significance. Annotated genes highlight candidates tested in *Xenopus* NC cells.

**Supplement Figure 12:**
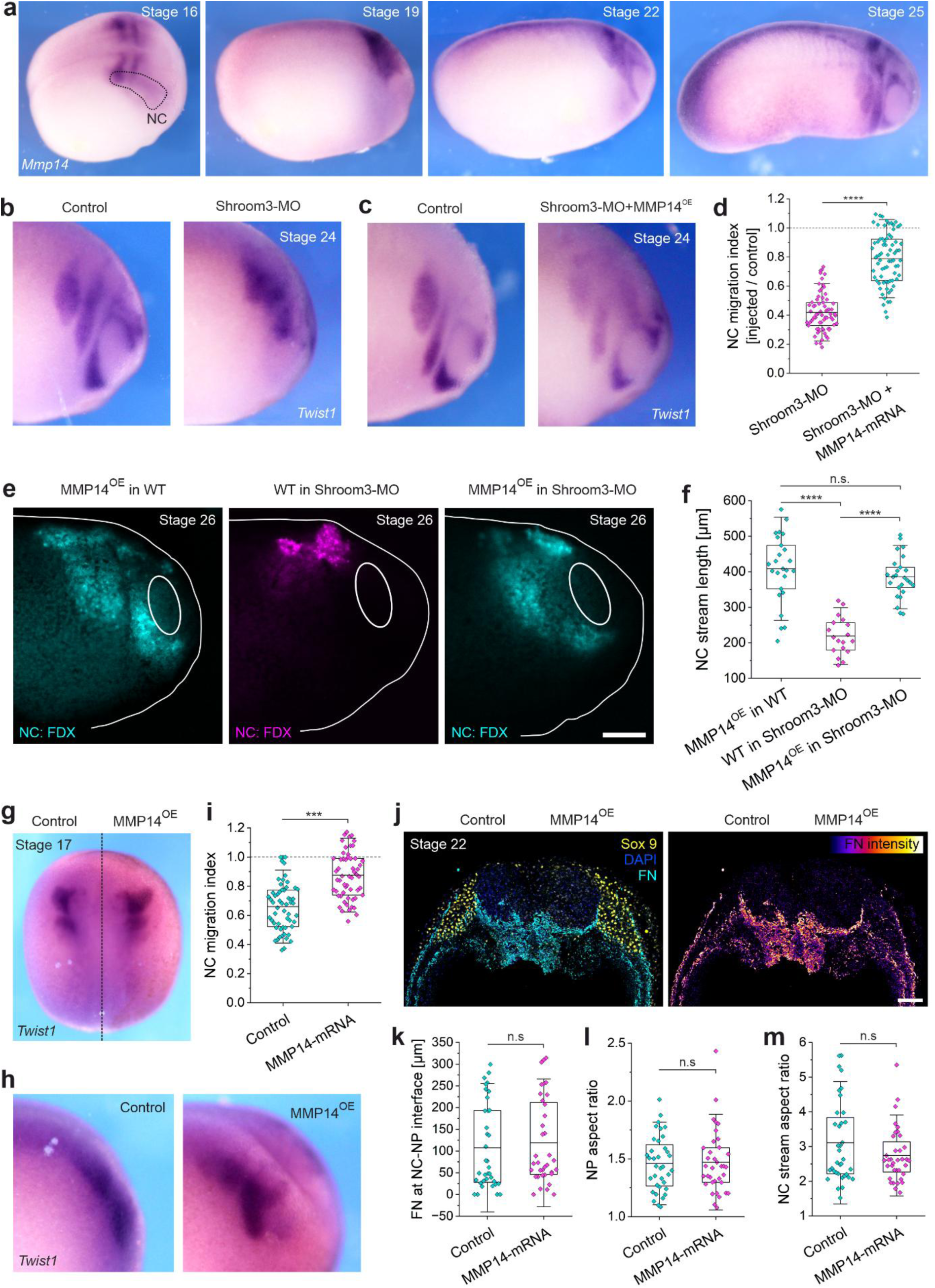
MMP14 control experiments in vivo. **a**, *Mmp14* expression monitored by ISH over stages during cephalic neurulation and NC migration. **b**-**c**, Embryos were injected with *Shroom3* morpholino alone (**b**) or together with mRNA encoding MMP14 (**c**) to achieve overexpression (MMP14^OE^). NC cells were stained for the NC marker *Twist1* by ISH. **d**, NC migration index in embryos injected as described (**b**-**c**). Index was calculated by dividing NC stream length in the injected and contralateral control side. **e**, Grafts of fluorescently labeled NC cells. MMP14^OE^ NC cells (cyan) were transplanted in either WT or Shroom3 morphant hosts (left and right panel), while WT NC cells (magenta) were transplanted into Shroom3 morphants (middle panel). **f**, Length of grafted NC streams at stage 26. Parts of data for the WT NC grafts in Shroom3-MO hosts are re-plotted in Supplement Figure 9b. **g**-**h**, NC migration in control (left embryo half) and MMP14^OE^ (right embryo half), analyzed by colorimetric ISH of the NC marker *Twist1*. **i**, NC migration index, calculated by normalizing NC stream length to the maximum value in the control. **j**, Transverse section of MMP14^OE^ embryo during NC migration (stage 22). Sections were stained for the NC marker Sox9 (yellow), FN (cyan and intensity LUT in right panel), and nuclei (DAPI). **k**-**m**, Quantifications of FN length at NC-NP interface (**k**), NP aspect ratio (**l**), and NC stream aspect ratio (**m**) in control and MMP14^OE^ embryo halves. Scale bars represent 200 µm (**e**) and 100 µm (**j**). For **d**, **f**, **i**, **k**, **l**, **m**, data are mean ± s.d. Statistical analysis was performed using two-tailed *t*-test (**d**, **i**), Tukey’s test (**f**), and two-tailed Mann-Whitney test (**k**-**m**); n.s. *P* > 0.05; \*\*\**P* ≤ 0.001; \*\*\*\**P* ≤ 0.0001. *n* = 69 Shroom3-MO and 73 Shroom3-MO + MMP14^OE^ embryos (**d**); *n* = 25, 18, and 25 grafts, respectively (**f**); *n* = 57 embryos (**i**); *n* = 36 sections from 10 embryos for each (**k**-**m**).

## Supplement Videos

**Supplement Video 1**: Dispersion and migration of control (left) and MMP14-depleted NC cells ex vivo.

**Supplement Video 2**: NC invasion into a FN matrix ex vivo. NC-mesoderm sandwich explant (as described in Figure 2i) with labelled NC cells (nuclear GFP) and labelled FN (Alexa Fluor 555).

**Supplement Video 3**: Retrograde FN displacement ex vivo. NC-mesoderm sandwich explant (as described in Figure 2i) with labelled NC cells (membrane GFP) and labelled FN (Alexa Fluor 555).

**Supplement Video 4**: Dorsal view on a Lifeact-RFP expressing embryo during neurulation. The right side of the embryos was injected with a morpholino against MMP14 to prevent NC-mediated FN remodeling.

**Supplement Video 5**: Orthogonal reslice (YZ) through cephalic NT region during neurulation. WT (cyan) and MMP14-MO embryos (magenta) express Lifeact-RFP in one embryo half. The dorsal midline is to the right side each.

**Supplement Video 6**: Scanning Electron Microscopy Array Tomography volume of NT region in a WT embryo, with fully segmented NT cell shape and nucleus.

**Supplement Video 7**: Scanning Electron Microscopy Array Tomography volume of NT region in a MMP14-MO embryo, with fully segmented NT cell shape and nucleus.

**Supplement Video 8**: Time-lapse movie of deep NT cells expressing mGFP in WT (left) and MMP14-MO (right). The superficial epidermis additionally expresses Lifeact-RFP. Individual cell shapes were outlined and projected for visualization.

**Supplement Video 9**: Time-lapse movie of mGFP (cyan) and Lifeact-RFP (magenta) expressing NP explants on Fibronectin (left) and Laminin (right).

**Supplement Video 10**: Dispersion and migration of NC explants from control (left) and Shroom3-depleted embryos ex vivo.

**Supplement Video 11**: Laser ablation of cell junctions in Lifeact-RFP expressing embryos at stage 15 and 20, and at stage 20, injected with a *Shroom3* morpholino or a CA-MYPT construct.

**Supplement Video 12**: Fluorescently labeled NC grafts, pre-compressed or uncompressed, and transplanted in WT or Shroom3 morphant hosts.

## Materials and Methods

### Animal procedures

*Xenopus laevis* embryos were obtained from a wild-type strain as described^[88]^. Ovulation of mature females (2-5 years of age) was induced by subcutaneously injecting 100 IU pregnant mare serum gonadotrophin (Intervet) into the dorsal lymph sac. A second injection of 200-300 IU human chorionic gonadotrophin (Intervet) was performed 72 h later. Eggs were fertilized *in vitro* by mixing with a sperm solution. Testes were provided by the European Xenopus Resource Centre in Portsmouth. Embryos were staged according to Nieuwkoop and Faber^[89]^. Fertilized eggs were dejellied in a 2 % (w/v) solution of L-cysteine (Sigma-Aldrich), neutralized with 500 µL 5 N NaOH, and maintained in 0.1× MMR (Marc’s Modified Ringer’s) or 3/8 NAM (normal amphibian media). Animal licenses were approved by the Animal Welfare and Ethical Review Board (WERB) at University College London and the UK Home Office. All animal experiments followed the relevant guidelines and regulations.

### Xenopus illustrations

Illustrations in Supplement Figures 5 and 9 are adapted from Normal Table of *Xenopus* development (Zahn drawings)^[89]^. Digital images created by Xenbase (http://www.xenbase.org/, RRID: SCR_003280)^[90]^.

### mRNA synthesis, morpholinos and microinjection

Capped mRNAs were transcribed from linearized plasmid templates using mMESSAGE mMACHINE SP6 or T7 kits (Thermo Fisher Scientific #AM1340 and #AM1344), according to manufacturer’s instructions. Linearized plasmids were purified using Monarch PCR & DNA Cleanup kit (New England Biolabs #T1130). Capped mRNA was purified using Monarch RNA Cleanup kit (New England Biolabs #T2040). Membrane GFP (mGFP)^[91]^, nuclear RFP (nRFP)^[91]^, Lifeact-RFP^[92]^, and Xsnail-N-GR^[38]^ (iDN-Snail) expression plasmids have been described previously. L_NLS 41kDa was a gift from Pere Roca-Cusachs (Addgene #201342; http://n2t.net/addgene:201342; RRID: Addgene_201342). CA-MYPT mCherry was a gift from Masazumi Tada (University College London). Plasmids with cDNA encoding for *Xenopus laevis Fn1* (MXL1736-202772769) and *Mmp14* (OXL11194-247068881) were purchased from the *Xenopus* Gene Collection at Horizon Discovery. The full-length *fn1.S* sequence was subcloned into a pCS2+ TA-eGFP backbone via an In-Fusion cloning kit (Takara Bio Inc.) to obtain pCS2-FN-eGFP.

Morpholinos used in this study were synthesized by Gene Tools LLC and prepared according to manufacturer’s instructions. Morpholinos targeting *Cxcr4^[37]^, Fn1^[50]^, Mmp14^[36]^, Shroom3^[23, 93]^, and Twist1^[94]^* were characterized and described previously.

A calibrated needle was used to microinject 5 nL solution into the following blastomeres to target the embryos’ right embryo half. *Fn1*-MOs or pCS2-FN-eGFP were injected at eight-cell stage, either into two animal blastomeres (animal-dorsal and animal-ventral) to target prospective ectodermal tissue or two vegetal blastomeres (vegetal-dorsal and vegetal-ventral) to target prospective mesodermal tissue. *Twist1*-MO, *Cxcr4*-MO, *Mmp14*-MO, *Shroom3*-MO, *Mmp14* mRNA, pCS2-FN-eGFP, L_NLS 41kDa, mGFP, nGFP, nRFP, and Lifeact-RFP were injected at eight-cell stage into two animal blastomeres. CA-MYPT mCherry was injected into 2 blastomeres at 16-cell stage (V1.1 & V1.2). iDN-Snail was injected into 1 blastomere at two-cell stage, and the construct was induced at stage 16 by adding 10 µM freshly dissolved Dexamethasone (Thermo Fisher Scientific #4902) in EtOH to 0.1× MMR. Mosaic expression of L_NLS 41kDa or pCS2-FN-eGFP was achieved by microinjecting circular plasmid DNA^[95]^. FITC- or Rhodamine-Dextran (Thermo Fisher Scientific #D1821 and #D1824) were used as fluorescent tracer to ensure correct blastomeres were targeted, if no fluorescent protein was co-injected. Microinjections were carried out in 3/8 NAM supplemented with 3% Polysucrose 400 (Sigma-Aldrich #P7798). 1°h after injection, embryos were maintained in 0.1× MMR.

Following concentrations were injected per blastomere: *Cxcr4*-MO: 8 ng; *Fn1*-MO: 20 ng each*; Mmp14*-MO: 7.5 ng; *Shroom3*-MO: 15 ng each*; Twist1*-MO: 1 ng; iDN-Snail: 0.75 ng; CA-MYPT mCherry: 0.8 ng; *Mmp14* mRNA: 0.5 ng; L_NLS 41kDa: 0.15 ng; FN-eGFP DNA: 0.15 ng; nRFP, nGFP, mGFP, and Lifeact-RFP: 0.3 ng each.

### Conjugation of FN antibody and labelling of endogenous FN

To label endogenous FN in living samples, an antibody against *Xenopus laevis* FN (clone 4H2, DSHB) was directly conjugated using Alexa Fluor 555 Antibody Labeling Kit (Thermo Fisher Scientific #A20187). Briefly, the antibody was purified from the supernatant using NAb Protein A/G Spin kit (Thermo Fisher Scientific #89950), desalted using Zeba Spin Desalting Columns with 7K MWCO (Thermo Fisher Scientific #89889), and subsequently concentrated using Pierce Protein Concentrators PES with 3K MWCO (Thermo Fisher Scientific #88514). The labeling reaction was carried out according to manufacturer’s instructions and excess dye was removed to purify the conjugated antibody. A total of 20 µL of conjugated antibody (with a stock concentration of 100 µg/mL) was distributed throughout the cephalic NC site of stage 17 embryos by four subcutaneous microinjections of 5 nL each. After 30 min incubation, the tissue was dissected and mounted for time-lapse recordings, as described in following sections.

### Biocompatible microdroplets

Biocompatible ferrofluid was a gift from Alessandro Mongera and used as previously described^[60]^. Briefly, 5 nL of the ferromagnetic fluid was injected subcutaneously into the region of the cephalic NC at the according stage using a calibrated needle. Magnetic tweezers were used to keep the microdroplet in place when retracting the needle and to manipulate its position, making sure the microdroplet was embedded between in the cephalic NC. Embryos were left to heal in NAM 3/8 for 30 min and subsequently fixed in 4 % PFA for 2 h at room temperature. After three consecutive washes in PBS, embryos were dehydrated in 100 % MeOH for 30 min with the MeOH being renewed every 10 min and subsequently cleared in BABB (1 part benzyl alcohol to 2 parts benzyl benzoate). Imaging was performed using a Leica SP8vis confocal microscope with a 10× dipping lens (HC APO L 10×/0,30W U-V-I).

Shapes of ferromagnetic droplets were quantified from maximum projections. To determine elongation and elongation angle of droplets, images were oriented along the anterior-posterior and dorso-ventral axis of the embryo, with anterior facing to the right (corresponding to 0°) and dorsal facing upwards (corresponding to 90°). Using FIJI^[96]^, droplet outlines were segmented by manual thresholding, an ellipse was fitted to the shape, and the aspect ratio of the ellipse as well as the angle was determined using the *fit ellipse* function in the measurement settings. The droplet aspect ratio was plotted against the elongation angle in a compass plot in OriginPro 2023 (https://www.originlab.com/), with the angle indicating the elongation direction of the droplet in respect to the embryo axis, and the length of the arrow indicating the elongation ratio itself.

### Laser ablation

Photoablation experiments were carried out using a Leica Stellaris 8 DIVE multiphoton microscope equipped with a 25× water dipping lens (HC IRAPO L 25×/1,00W motCORR) and a dual-beam Coherent Discovery NX laser. The laser wavelength was tuned to 920 nm and set to 16% power. The experiment was designed using the FRAP wizard on LAS X software. After acquiring three default images, junctions were ablated on a small, manually defined ROI and on a single z plane with 3 iterations. Junction recoil was subsequently monitored for 20 seconds. No more than 3 ablations were performed per embryo, and embryos were screened for photo damage after every ablation experiment. To calculate recoil velocity, recoil distance was tracked from the left and right vertex to the respective cutting edge for 10 s post ablation using the Manual Tracking plugin in FIJI.

### Compression in vivo and ex vivo

Exogeneous pressure was applied on cephalic NC in vivo using a custom-built device. The tip of a borosilicate glass capillary (WPI #1B100F-4) was cut and fire polished to prevent damage to the embryo. Stage 16 embryos were mounted in molding clay and the blunt tip placed onto the embryo, targeting the ventral region adjacent to the NC front. The needle was held in place by a micromanipulator for 7 h.

Exogeneous pressure was applied on cephalic NC explants by compressing stage 16 NC clusters between the plastic dish and a glass coverslip that was held in place by high-vacuum grease (Sigma-Aldrich #Z273554). Compression was applied by gently squeezing the coverslip with forceps until the NC clusters visibly flattened without destroying the cluster integrity, and the compression was held in place for 4 h, corresponding with the timing of NT closure between stage 16 and 20 at 22°C. NC clusters were subsequently used as grafts in transplantation experiments or lysed for RNA extraction and qPCR analysis.

### Transplantation (grafting) experiments

Cephalic NC was dissected using a custom-made hair knife as previously described^[97]^. The overlying superficial ectoderm was lifted off and the endogenous NC removed. Dissection was performed in 3/8 NAM. Grafting was also performed as previously described^[98]^. The cephalic NC was removed and either transplanted directly or cultivated for 4 h ex vivo under compression or uncompressed in 1× DFA (Danilchik’s For Amy) before being transplanted to the region immediately dorsal to the cranial placodes. The overlying ectoderm was unfolded back over the NC, and a glass coverslip was placed on top until the embryo fully healed, after which the coverslip was removed.

### Explant cultures

Cephalic NP explants were dissected from stage 14 embryos with a hair knife. The most superficial layer of the neuroepithelium was removed between the emerging neural folds and small clusters of NP cells were lifted off, making sure to explant NP tissue only. To promote adhesion, explants were chopped into smaller pieces using a hair knife and transferred to an FN- or LN-coated dish, filled with 1× DFA. For ex vivo intercalation assays, two NP explants, one expressing mGFP and one Lifeact-RFP, were placed in proximity, using a hair knife to orient explants.

Cephalic NC explants were dissected from stage 16 embryos. Using a hair knife, the superficial epidermis was removed and the NC lift off, as previously described^[97]^. Similarly, cephalic NC explants were dissected from stage 20 embryos, 30 min after the Alexa Fluor 555-conjugated FN antibody was injected subcutaneously, to explant fragments of endogenous FN together with the cephalic NC.

NC-mesoderm sandwich explants were dissected from stage 16 embryos, 30 min after the Alexa Fluor 555-conjugated FN antibody was injected subcutaneously. The superficial epidermis was removed above the cephalic NC using a hair knife and the NC was separated from the NP by making an incision dorsally along the NC-NP interface. Three more incisions were made, one ventral and two lateral from the cephalic NC, to mark the edges of the explant. The hair knife was subsequently inserted underneath the mesoderm to scrape off the connection to the endoderm and lift off the mesoderm/cephalic NC explant. The explants were transferred to an FN-coated dish, oriented with the NC facing upwards, and immediately sandwiched, using a second FN-coated coverslip that was held in place by high-vacuum grease (Sigma-Aldrich #Z273554).

Substrates were coated by adsorbing 100 µg mL^-1^ FN (Sigma-Aldrich #F1056) or LN (STEMCELL Technologies #77003) to the glass-bottom dishes or coverslips overnight before the experiment at 4°C.

### Cryosectioning

Embryos were fixed in 4% PFA overnight at 4°C, washed three consecutive times in PBS for 5 min, and dehydrated in 30% sucrose in PBS overnight at 4°C. Embryos were then embedded in a 1:1 solution of 30% sucrose in PBS and OCT mounting media (VWR #361603E), frozen on dry ice and cryosectioned into 30 µm slices. Slides were incubated at 37°C for 1 h and then stored at room temperature overnight, before proceeding with subsequent immunostaining.

### Electron microscopy sample preparation and Scanning Electron Microscopy Array Tomography (SEMT-AT)

Whole *Mmp14*-MO embryos were fixed with EM grade 2% formaldehyde / 1.5% glutaraldehyde in 0.1M sodium cacodylate overnight at 4°C. The following day, embryos were incubated with 1% osmium tetroxide / 1.5% potassium ferricyanide for 1 hour at 4°C, 1% thiocarbohydrazide for 20 min, 2% osmium tetroxide for 30 min and 1% uranyl acetate overnight, with several ddH_2_O washes between each incubation. Samples were then dehydrated through an ethanol series and infiltrated with propylene oxide:Epon (50:50, 1 hour) then two 100% Epon incubations (>2 hours each) prior to oven polymerization at 60°C. Semithin and ultrathin sections were collected and screened by light and transmission electron microscopy respectively to target transverse cross sections of the NP. Once targeted in the appropriate orientation, serial ultrathin sections (∼500 sections) were collected onto ITO coated coverslips manually using the paperclip method^[45]^ and screened by backscattered electron imaging at 100 nm px to identify NC cells that were fully encompassed within the 500 serial section array. ROIs capturing NC cells in both control and morpholino-injected sides were imaged at 10 nm px using the Sense backscatter detector (Zeiss), with an accelerating voltage of 4.5kV, stage deceleration at 3 kV and landing energy of 1.5 kV using a Gemini 300 (Zeiss) with Atlas 5 (Fibics). Images were aligned with TrakEM2^[99]^ in FIJI^[96]^, and cell and nuclear boundaries were segmented, 3D reconstructed and movies created using Amira software^[100]^ (ThermoFisher Scientific).

### In situ hybridization

Colorimetric whole-mount in situ hybridization was performed as previously described^[98]^. In brief, *Adam13*-, *Fn1*-, *Mmp2*-, *Mmp14*-, and *Twist1*-digoxigenin riboprobes were transcribed using the Riboprobe in vitro Transcription System (Promega #P1420). The plasmid containing the *Twist1* probe was cloned in the lab, the *Adam13* probe was gifted by Dominique Alfandari (University of Massachusetts Amherst), the *Mmp2* probe was a gift from Yun-Bo Shi (National Institutes of Health). cDNAs encoding transcripts of *Xenopus laevis Fn1* (MXL1736-202772769) and *Mmp14* (OXL11194-247068881) were purchased from Horizon Discovery. *Mmp14* probe was generated by linearizing the plasmid using BamHI restriction and T7 in vitro transcription. For the *Fn1* probe, the 0.8 kB c-terminal fragment was amplified via PCR and ligated into pGEM^®^-T Easy (Promega #A1360) for SP6 in vitro transcription.

Embryos were fixed in MEMFA overnight at 4°C, bleached in 6 % hydrogen peroxide bleaching buffer, and then incubated with the according probes overnight in hybridization buffer. Embryos were then washed, blocked with 2 % blocking reagent (Sigma-Aldrich #11096176001), incubated with 1:2,000 anti-digoxigenin-AP antibodies (Sigma-Aldrich #11093274910), and then revealed using NBT/BCIP with AP buffer. Embryos were imaged using the Nikon SMZ800N attached to the DS-Fi3 camera (Nikon DS-L4).

Fluorescent in situ hybridization was performed as previously described^[101]^. In brief, MEMFA-fixed embryos were incubated with the *mmp14* probe overnight in hybridization buffer. Embryos were then washed, bleached in 3 % hydrogen peroxide, and incubated with 1:1,000 anti-digoxigenin-POD antibody (Sigma-Aldrich #11207733910). After washing, embryos underwent the fluorescent POD reaction with the Cy5-tyramide solution (Quanterix #NEL745001KT). For subsequent immunostaining, embryos were cryosectioned and stained as described.

Hybridization chain reaction (HCR)^[102]^ on cryosection was performed using a modified protocol from Molecular Instruments. DNA probes against *Xenopus laevis Fn1* were designed by Chintan Trivedi (University College London) using a custom-built software tool^[103]^ (https://github.com/ctucl/pybridizer). Hairpins, buffers, and wash solutions were purchased from Molecular Instruments (https://www.molecularinstruments.com/shop). Embryos were fixed in MEMFA overnight at 4°C, washed in PBS, and dehydrated in an EtOH series. 0.4 pmol of each probe was diluted in probe hybridization buffer and the sections were incubated overnight at 37°C. Slides were washed in pre-warmed wash solution and excess wash buffer was removed in a wash series of 5× SSCT. For amplification, slides were pre-amplified in a humidified chamber for 30 min at room temperature, while 15 pmol of hairpin h1 and h2 each were assembled at 95°C and snap cooled to room temperature. Hairpins were added to amplification buffer and slides were incubated overnight at room temperature. Excess hairpins were washed off with 5× SSCT. For subsequent immunostaining, sections were additionally treated as described in the following.

### Immunostaining

Cryosections were permeabilized with 1×PBS+0.1 % TritonX-100 for 30 min, blocked in 5 % BSA for 1 h and then incubated in primary antibodies, diluted in 1 % BSA, overnight at 4°C. Primary antibody solutions were removed in three consecutive wash steps with PBS + 0.1 % Tween20, and the slides were then incubated in secondary antibodies and affinity probes, diluted in 1 % BSA for 2 h at room temperature. Sections were washed in PBS + 0.1% Tween20 and embedded in Fluoromount-G (Thermo Fisher Scientific #00-4958-02).

Whole mount immunostaining of *Xenopus laevis* embryos was performed in a similar way but with extended incubation of secondary antibodies/affinity probes (overnight at 4°C) and washing periods (5× 20 min in PBS+0.1 % Tween20).

NP explants were fixed for 25 min in 4 % PFA at room temperature, permeabilized with 1× PBS + 0.1 % TritonX-100 for 20 min, and then incubated with primary antibodies, diluted in 1 % BSA, for 1 h at room temperature. Primary antibodies were removed by washing with PBS + 0.1 % Tween20, and explants were incubated with secondary antibodies and affinity probes, diluted in 1 % BSA, for 1 h at room temperature. Excess antibodies were washed off with PBS + 0.1 % Tween20 and explants were stored in PBS.

Following primary antibodies and dilutions were used: mouse α-fibronectin (clone 4H2, DSHB) at 1:100; goat α-Sox2 (R&D #AF2018) at 1:200; rabbit α-Sox9 (Sigma-Aldrich #AB5535) at 1:500; mouse α-GFP (Thermo Fisher Scientific #A-11120) at 1:100; rabbit α-GFP (Thermo Fisher Scientific #A-11122) at 1:100; rabbit α-Laminin (Sigma-Aldrich #L9393) at 1:100; rabbit α-Rab11A (Thermo Fisher Scientific #700184) at 1:200; mouse α-aPKC (Santa Cruz #sc-17781) at 1:100; rabbit α-Phospho-Myosin Light Chain 2 (pMLC) (Cell Signalling Technology #3671) at 1:100.

Secondary antibodies conjugated with Alexa Fluor 488, 568, or 647 were purchased from Thermo Fisher Scientific and diluted 1:250; DAPI (Thermo Fisher Scientific #62248) was diluted 1:1000; Phalloidin conjugated with Alexa Fluor 568 or 647 (Thermo Fisher Scientific #R415 and #A22287) was diluted 1:250.

### Human induced pluripotent stem cells (hiPSC) and induced neural crest cells (iNCC)

iNCCs were generated from healthy hiPSC and characterized as previously described^[104]^. iPSCs used in this study were NIBSC8, obtained from the National Institute for Biological Standards and Control (UK). Briefly, iPSCs grown in E8 media (Thermo Fisher Scientific # A1517001) were dissociated into single cells using TrypLE Select (Thermo Fisher Scientific A1285901) and plated at 20.000 cells/cm^2^ in Neural Crest Medium (DMEM/F12, 0,5 % BSA, 1× GlutaMAX, 1× B27 Supplement and 3 µM CHIR99021) for five days. Both hiPSCs and iNCCs were grown in Geltrex (Thermo Fisher Scientific #A4000046703) coated dishes.

Cell compression was conducted using Transwell migration assays in 6-well plates with 8 µm pore ThinCerts (Greiner Bio-One #657638). A total of 5000 iNCCs were plated on Geltrex-coated inserts in NCC medium without CHIR99021. After 24 h, cells that migrated through the pores to the bottom compartment (compressed fraction) and cells remaining on the upper surface of the insert (uncompressed fraction) were harvested separately by enzymatic dissociation using TrypLE Select.

### RNA isolation and sequencing

RNA was extracted from cell fractions using the Monarch Total RNA Prep Kit (New England Biolabs #T2110), following manufacturer’s instructions. RNA quality was assessed by NanoDrop measurements and Tapestation fragment analysis. RNA samples (three each, compressed and uncompressed) were sequenced by BGI Genomics using proprietary library prep kits with poly(A) enrichment and DNBSeq technology. FASTQ files containing approximately 50 million reads per sample were obtained and used for downstream differential expression analysis.

We used the Cactus pipeline (https://github.com/jsalignon/cactus) for differential expression analysis. In short, Cactus uses FastQC and MultiQC for pre-processing and quality control, followed by quantification of raw reads using pseudoalignment with kallisto against the human reference transcriptome (hg38). Transcripts and genes are analyzed for differential expression using sleuth, and false discovery rates (FDRs) are computed.

### Quantitative PCR (qPCR) analysis of gene expression

RNA was extracted from a pool of 25 cephalic NC explants (compressed or uncompressed controls) using TRIzol reagent (Thermo Fisher Scientific #15596026) according to manufacturer’s instructions. qPCR reaction mix was set up using the Luna Universal One-Step RT-qPCR kit (New England Biolabs #E3005L) following manufacturer’s instructions and with a total of 8 ng RNA for each sample. Reactions were performed in a QuantStudio 3 qPCR System (Thermo Fisher Scientific) using standard parameters with an initial step for reverse transcription (55°C for 10 min). Relative expression values (Delta Ct) were calculated as previously reported^[105]^, using *Ef1*α and *Odc1* as endogenous controls. Primers for targets in this study were designed in exon-exon junctions using the Primer Blast tool (NCBI).

*Mmp14* gene expression quantification was achieved using multiplex probe-based qPCR. *Mmp14* and *Odc1* primers and probes were designed using Primer quest online tool (IDT) and ordered through Merck. *Mmp14* probe had 5’ 6-FAM and 3’ BHQ1 modifications. *Odc1* probe, used as reference housekeeping gene, had 5’ ROX and 3’ BHQ2 modifications. One-step qPCR was performed using 2 ng of total RNA, 250 nM and 900 nM of each probe and primers, respectively, in a TaqPath One Step Multiplex master mix reaction, following manufacturer’s recommendations. qPCR reactions were run in a BioRad CFX 96 real-time PCR system and Ct values were obtained for relative expression quantification. Relative expression values were calculated using the 2^Delta Ct method, in which Cts of test samples were subtracted from Cts of control samples. *Odc1* values were used to calculate a normalization factor by the same method and used as a dividing factor for *Mmp14* values.

Primer sequences used in this study are summarized in Table 1.

**Table 1:**
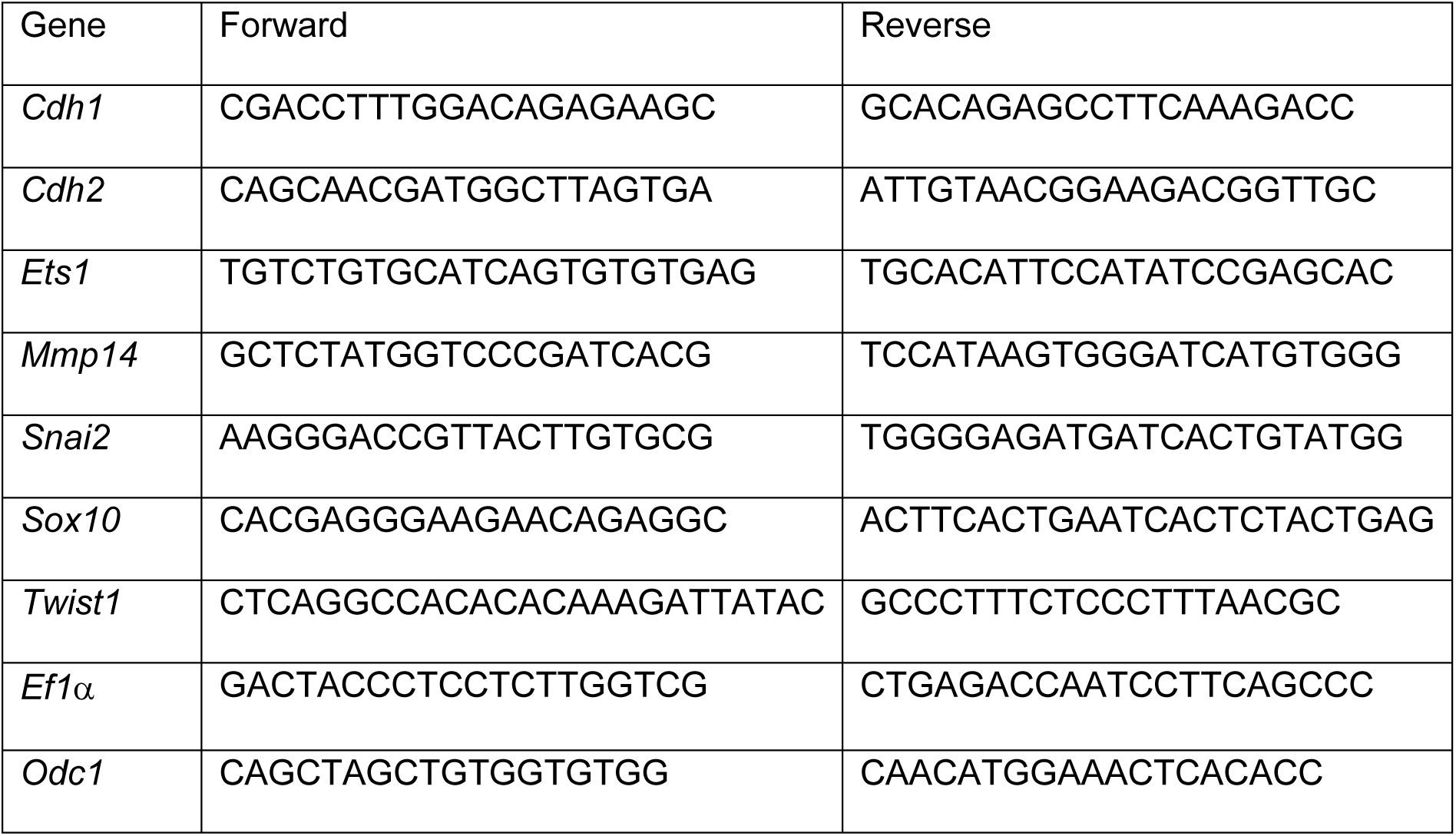
Primer sequences used in this study.

### Microscopy and image treatment

Confocal stacks were acquired on a Leica SP8vis, using 10× (HC PL FLUOTAR 10x/0,30) or 20× (HC PL FLUOTAR 20x/0,50) magnification. High resolution confocal stacks were acquired on a Zeiss LSM 980, operating in the 4Y Airyscan mode and using 25× magnification with water immersion (LD LCI Plan-Apochromat 25×/0.8 Imm autocorr DIC M27). NC grafts were imaged on a Leica M205 FCA fluorescence stereo microscope.

Time-lapse recordings of whole embryos, NC grafts, and NC-mesoderm sandwich explants were acquired on a Leica SP8vis, using 10× (HC APO L 10×/0,30W U-V-I) or 20× (HC APO L 20×/0,50 W U-V-I) dipping lenses. Brightfield time-lapse recordings of WT, MMP14-MO, and Shroom3-MO NC dispersion as well as epifluorescence time-lapse recordings of intercalating NP explants were acquired on a compound microscope, equipped with a 10× lens (Achrostigmat 10×/0.25 Ph1) and Simple PCI v.6.6 software.

General treatments including maximum intensity projection, contrast adjustment, substacking, image rotation, resizing, reslicing, color-coded projection, pseudocoloring, overlaying, scaling, and annotation were done in FIJI^[96]^. All adjustments were made to enhance clarity without interfering with fluorescence intensity values.

Color-coded projections in Figure 3e were done by reslicing the respective WT or MMP14-MO time-lapse image stack in YZ plane, sub-stacking a volume of 25 Z slices along the ROI, maximum projecting the substack, and color-coding individual frames. Cell shapes were segmented by freehand selection using the mGFP signal and the resulting shape outlines were temporally aligned in a stack and projected as temporal color-code.

### Quantification and data analysis

Quantification of FN length and intensity along interfaces was done on maximum intensity projections of immunostained cryosections by manually drawing segmented lines along the ROI in FIJI. Background intensities were subtracted from individual projections, when calculating the sum of NC interfaces.

*Fn1* and *Mmp14* mRNA intensities were quantified from HCR or fluorescent ISH staining in cryosections, respectively. ROIs were selected manually based on immunofluorescent co-staining of Sox9 and FN.

NT area and aspect ratios were determined by segmenting freehand selections along the ROI. Overlay plots of NT hemispheres or NC streams were done by aligning and overlaying individual transparent projections in CorelDRAW. NT closure index was calculated by dividing NT hemisphere areas in the treated versus the control embryo half.

NC and NT density was calculated from cryosections by counting nuclei in the ROI and dividing the number of nuclei by the ROI area, as determined by segmented freehand selections of the tissue area. Aspect ratios of cells and nuclei in Figure 3j and Supplement Figure 7d were determined by segmenting individual shapes in the highlighted ROIs of SEM sections.

N/C ratio of L_NLS 41kDa was determined as described^[61]^ and quantified in maximum intensity projections of individual NC cells. The sensor signal in Sox9-positive nuclei was segmented using DAPI staining as a mask, and the cell shape (without the nucleus) was segmented using the GFP signal. N/C ratio was calculated as:

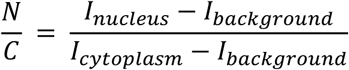

With *I_nucleus_* and *I_cytoplasm_* being the respective mean fluorescence intensities and *I_background_* the mean intensity of a cell negative for GFP.

Apical constriction of NT cells was quantified by segmenting cell surfaces in Cellpose^[106]^ (https://www.github.com/mouseland/cellpose). The mean surface area of all segmented NT cells from WT and MMP14-MO embryos was plotted as a function of time, with the apical area measured at each time point normalized to time zero.

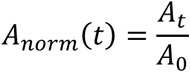

For each cell, the mean rate of change in normalized apical area over 8 hours was used as a summary measure of apical-area dynamics:

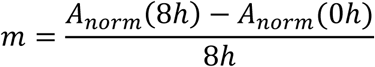

NC stream length in colorimetric ISH images was determined by measuring the length of the second stream in the treated and control side. NC stream length of grafts was measured similarly but shown as absolute values due to a missing internal control. NC migration index was calculated by dividing NC stream length of treated versus control embryo halves, illustrated in Supplement Figure 3g, or by normalizing to the maximum value in the control. NC migration index in cryosections was calculated by dividing length of FN at the NC-ectoderm interface in the treated versus control embryo half.

Analysis of NC explants ex vivo was done by manually tracking NC cells in FIJI. Directionality and velocity were calculated from these tracks using the Chemotaxis and Migration Tool (Ibidi). The full formulas for calculation can be found on the Ibidi website (https://ibidi.com/chemotaxis-analysis/171-chemotaxis-and-migration-tool.html).

NP intercalation ex vivo was analyzed in FIJI. After thresholding, binary masks were generated for tissue areas, and the overlap area occupied by both binary masks was determined. An overlap index was calculated as:

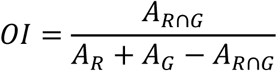

Actin intensity of individual NP clusters was measured on fixed samples using the phalloidin staining and line scans along individual junctions. Height and spreading area of individual NP clusters was analyzed in axial stacks and by manually segmenting the area in contact with the adhesive substrate, respectively.

### Statistics and reproducibility

Normality in the spread of data for each experiment was tested using Shapiro-Wilk, Kolmogorov–Smirnov, Lilliefors, Anderson-Darling, d’Agostino–Pearson and Chen-Shapiro tests. Significances for datasets displaying normal distributions were calculated with an unpaired two-tailed Student’s t-test or one-way analysis of variance with post hoc Dunnett’s or Tukey’s test for multiple comparisons. Significances for non-normal distributed data were calculated using a two-tailed Mann-Whitney-Tests or Kruskal–Wallis tests with post hoc Dunn’s correction for multiple comparisons. All tests were calculated in OriginPro 2023. Asterisks of statistical significance or lack of statistical significance refer to the following probability values: n.s *P* > 0.05; \**P*≤ 0.05; \*\**P* ≤ 0.01; \*\*\**P* ≤ 0.001; \*\*\*\**P* ≤ 0.0001.

Apart from array tomography data, all experiments were performed at least three times. Note that for in vivo compression (Figure 5h&i), due to access to two compression devices only, individual experiments were performed with one embryo, repeated over 8-9 times.

Note that some data are re-plotted due to multiplexing of experiments and resourceful use of embryos. For NC migration data in Supplement Figure 4k&l and Supplement Figure 9e&f, parts of WT data serve as control for both Shroom3-MO and MMP14-MO. Multiplexed data for NC grafts in Shroom3-MO hosts are re-plotted in Supplement Figure 9b and Supplement Figure 12f. All re-plotted data are explicitly highlighted in the figure captions.

Wherever cryosections were used for quantitative data analysis, measurements were taken from sections of at least 10 different embryos (pooled from three independent experiments) and from several sections in each embryo along the ROI defined in Supplement Figure 1a.

## Acknowledgements

We thank Alessandro Mongera for kindly providing the ferromagnetic fluid, Ines Fernandez for helping with segmentation, William Andrews for frequent access to his Cryostat, Chintan Trivedi for designing HCR probes, and Masa Tada for gifting the CA-MYPT mCherry plasmid. We are also very thankful to Jiawen Peng and Nicole Baxter for their support with experimental procedures and data analysis. Work in the laboratory of R.M. is supported by grants from the Medical Research Council (MR/S007792/1), Biotechnology and Biological Services Research Council (M008517; BB/T013044), and Wellcome Trust (102489/Z/13/Z). The UCL, LMCB EM facility (RRID:SCR_027340) is supported by funding from the Wellcome Trust (218278/Z/19/Z). K.W. was supported by the Deutsche Forschungsgemeinschaft (DFG) through a Walter Benjamin fellowship (513518868), in addition to the R.M. grants above.

## Abbreviations

ECM: extracellular matrix
EMT: epithelial-to-mesenchymal transition
FN: fibronectin
LN: laminin
NC: neural crest
NP: neural plate
NT: neural tube

